# Lung Epithelial Regulation of BCL2 Related Protein A1 (BCL2A1) by Coronaviruses (SARS-CoV) and Type I Interferon Signaling

**DOI:** 10.1101/2021.07.21.453244

**Authors:** Chilakamarti V. Ramana

**Affiliations:** Dartmouth-Hitchcock Medical Center, Dartmouth Med School, Lebanon, New Hampshire

## Abstract

Highly pathogenic respiratory viruses such as 1918 influenza (HIN1) and coronavirus (SARS-CoV-2) induce significant lung injury with diffuse alveolar damage, capillary leak, and extensive cell death resulting in acute respiratory distress syndrome (ARDS). Direct effects of the virus, as well as host immune response such as proinflammatory cytokine production, contribute to programmed cell death or apoptosis. Alveolar lung epithelial type II (AT2) cells play a major role in the clearance of respiratory viruses, secretion of surfactant proteins and antimicrobial substances into the bronchoalveolar fluid as well as repair of lung injury. Gene expression in AT2 cells is regulated in a tissue and cell-specific manner and in a temporal fashion. The availability of tissue and cell-specific RNA datasets in Human Protein Atlas led to the identification of localized expression patterns of BCL-2 family members such as BCL2 related protein A1 (BCL2A1) in AT2 cells and immune cells of the lung. BCL2A1 expression was regulated by multiple stimuli including Toll-like receptor (TLR) ligands, interferons (IFNs), inflammatory cytokines, and inhibited by the steroid dexamethasone. In this study, regulation of BCL2A1 gene expression in human lung epithelial cells by several respiratory viruses and type I interferon signaling was investigated. SARS-CoV-2 infection significantly induced BCL2A1 expression in human lung epithelial cells within 24 hours that required the expression of Angiotensin-converting enzyme 2 (ACE2). BCL2A1 mRNA induction by SARS-CoV-2 was correlated with the induced expression of IFN-β and IFN-regulated transcription factor mRNA. BCL2A1 was induced by IFN-β treatment or by infection with influenza virus lacking the non-structural protein1(NS1) in NHBE cells. Furthermore, bioinformatics revealed that a subset of BCL-2 family members involved in the control of apoptosis and transcription such as BCL2A1, BCL2L14, BCL3, and BCL6 were regulated in the lung epithelial cells by coronaviruses and in the lung tissue samples of COVID-19 patients. Transcriptomic data also suggested that these genes were differentially regulated by the steroid drug dexamethasone.

## 1 Introduction

The alveolar epithelium of the human lung functions as a frontline cellular barrier for respiratory pathogens and is actively involved in the clearance of respiratory viruses, including coronaviruses (1, 2). There are two functionally distinct alveolar epithelial cells known as type I and type II (AT1 and AT2) representing major and minor cell populations, respectively (3, 4). AT2 cells are susceptible to SARS-CoV-2 infection as they co-express viral entry factors ACE2 and Transmembrane protein serine protease or TMPRSS2 (5). Lung type II cells provide a protective function to the lung by detoxification of pollutants and secretion of a variety of anti-inflammatory and antimicrobial substances into the alveolar fluid (6–8). In addition, lung type II cells can proliferate and undergo trans-differentiation into type I cells in response to lung injury (9) Significant lung injury frequently accompanies a highly pathogenic respiratory virus infection, which is mediated both by the direct effects of the virus as well as a result of the host immune response (10). Lung resident immune cells such as dendritic cells and natural killer (NK) cells produce significant amounts of antiviral and proinflammatory cytokines in response to virus infection, including type I and type II interferon (IFN), tumor necrosis factor-alpha (TNF-α), interleukin-beta 1(Il-1β) and interleukin 6 (IL-6) in a toll-like receptor (TLR)-dependent manner (11, 12).

Detection of proinflammatory cytokines in the bronchoalveolar lavage fluid (BALF) in COVID-19 patients was reported (13). Furthermore, bacterial products such as lipopolysaccharide (LPS) from secondary bacterial infection were also detected in the blood of COVID-19 patients (14). These proinflammatory cytokines and ligands activate multiple intracellular signal transduction pathways in alveolar epithelial cells resulting in the secretion of chemokines involved in the inflammatory influx of neutrophils, macrophages, lymphocytes, and viral clearance (15).

Cytokine mix of TNF-α, IL-1β, and IFN-γ is a potent inducer of cell death or apoptosis (16). Elimination of damaged or unwanted cells in biological systems is accomplished through apoptosis. These are genetically conserved and highly ordered responses critical during animal development and necessary to maintain homeostasis (17). Apoptosis can be morphologically distinguished by the presence of membrane blebbing, chromatin condensation, and DNA fragmentation. Apoptosis utilizes proteases known as caspases by two major pathways to initiate and execute cell death (18) The intrinsic pathway was activated by stimuli like DNA damage, nutrient starvation, and growth factor withdrawal leading to the loss of membrane polarization and release of cytochrome c from mitochondria. BCL-2 family members include anti-apoptotic (BCL2, BAK1) and proapoptotic (BAX, BID) members that play a major role in the regulation of apoptosis (19). In contrast, the extrinsic pathway is activated by the binding of cell death ligands like TNF-α or Fas ligand (FASL) to their corresponding receptors. Apoptosis was observed in AT1, AT2, endothelial cells, macrophages, and T cells in the autopsied lung sections of COVID-19 patients (20). Furthermore, both the intrinsic and extrinsic pathways of apoptosis were observed. Similar results were also obtained in the non-human primate (NHP) model of SARS-CoV-2 infection (20). Loss of cellular functions by apoptosis is a serious and frequent complication in patients suffering from microbial infections, multiorgan failure (MOF), and septic shock. Similarities between septic shock and COVID-19 including a dramatic increase in cytokine and chemokine levels, and an increase in apoptosis were noted (21). Diverse bacterial species and high levels of inflammatory cytokines TNF-α, IL-1β, and IFN-γ in the serum were demonstrated in septic shock patients (22). The role of the septic sera, and bacterial RNA products capable of activating Nuclear factor kappaB (NF-KB), Signal transducer and activator of transcription (STAT1), and Interferon regulatory factor 1 (IRF-1) in apoptosis was studied in the cell culture model of septic shock (22, 23). Requirement of components of interferon signaling such as Jak kinases, STAT1, IRF1, double-stranded RNA protein kinase (PKR), and caspases in sepsis and bacterial RNA mediated apoptosis was demonstrated (22–25).

BCL-2 family members are involved in the regulation of inflammation and apoptosis and play a major role in the growth, differentiation, and cancer of hematopoietic cells (26). However, regulation of inflammation and apoptosis in the lung AT2 cells by BCL-2 members remains to be investigated. Gene expression profiling in AT2 cells led to the identification of genes regulated by respiratory viruses and type I interferons (27). In this report, lung cell atlas RNA profiling led to the identification of distinct expression patterns in AT2 and immune cells of several BCL-2 family members. Furthermore, BCL2A1 and BCL2L14 were regulated by SARS CoV and type I interferons in lung epithelial cells and in the tissue samples of COVID-19 patients.

## 2 Materials and Methods

### 2.1 Gene expression datasets and Bioinformatics

Human and mouse lung type II- specific gene and cell atlas data were downloaded from Human Protein Atlas and LUNGGENS websites, respectively. Gene expression in human lung cell lines infected with respiratory viruses and COVID-19 patients was reported previously (28). Immgen RNA seq SKYLINE resources were used (http://rstats.immgen.org/Skyline_COVID-19/skyline.html). Microarray data in GEO datasets (GSE147507) (GSE47960) (GSE156295) was retrieved.. Supplementary data and gene expression datasets were also downloaded from the Journal publisher’s websites, Immgen browsers, and Signaling Pathway Project. Geo datasets were analyzed with the GeoR2R method (NCBI). Corresponding mock and treatment datasets were used. Outliers of expression were excluded in the data analysis. Cluster analysis was performed using gene expression software tools (29). Protein-protein interactions were visualized in the Search Tool for the Retrieval of Interacting Genes/Proteins (STRING) database (30). Gene ontology (GO), signaling and transcription factor analysis was done in KEGG. Metascape and TRANSFAC databases, respectively (31–33). Interferon-related data was retrieved from www.Interferome.org. Gene-specific information was retrieved from standard bioinformatics websites as described previously (27).

## 3. Results and Discussion

### 3.1 Profiling lung epithelial type II cell-specific expression of BCL2 family members involved in inflammation and apoptosis

Host-pathogen interactions play a major role in apoptosis resulting in the killing of the virus-infected cells or leading to enhanced viral replication and release (34). Mitochondria are known as the powerhouses of the cell due to their role in energy production. Furthermore, they are involved in antiviral responses and apoptosis. Mitochondrial antiviral-signaling protein (MAVS) coordinates innate immunity and type I interferon signaling to respiratory viruses (35). Some members of the BCL-2 family members such as anti-apoptotic BCL2 and pro-apoptotic BAX are located in the mitochondria and are involved in the control of apoptosis (19). In contrast, other members such as BCL2A1 and BCL6 respond to multiple inflammatory stimuli including TLR ligands, TNF-α, IL1-β, and antiinflammatory DEX and are implicated in the regulation of transcription, inflammation and apoptosis (36, 37). The selection of BCL-2 family members that respond to multiple inflammatory stimuli and regulated by DEX for cell atlas profiling was accomplished by using microarray datasets (Supplementary data). The intracellular location and function of the selected genes BCL2A1, BCL2L14, BCL11A, BCL3, and BCL6 were described (Table 1). The human lung is involved in the essential function of respiration and is composed of more than 30 cell types including epithelial endothelial, fibroblast as well as immune cells such as macrophages, granulocytes, and T cells. Human Protein Atlas (HPA) database is a useful resource providing RNA expression levels of genes in each cell type of an organ (38). Cell-specific RNA expression profiling in the human lung tissue of the HPA revealed that the selected BCL2 family members exhibit distinct expression patterns in immune cells and AT2 cells. For example, surfactant protein C (SFTPC) RNA expression was limited to C1 and the C6 AT2 cell populations in cell atlas, also known as the UMAP. In contrast, BCL2A1 RNA expression was predominantly located in the AT2 C6 and immune cell populations (Figure 1). Quantitative analysis revealed that BCL2 A1 RNA was highly expressed in the order of mac C2, mac C0, AT2 C6, and the granulocyte C4 cell populations. In contrast, BCL2L14 was highly expressed in the order of AT2 C6, club C7, ciliated C8, and the mac C2 cell populations (Figure 2). Heatmap representation of gene expression provides a comprehensive and convenient method of data visualization (39). Heat map representation of the selected BCL2 family members showed high expression levels in lung AT2 cells and the immune cells. AT2 cell-specific markers such as SFTPA2, SFTPC, LAMP3, and the AT1cell-specific markers such as AGER, CAV, and EMP2 were shown for a comparison purpose (Figure 3).

**Table 1.**
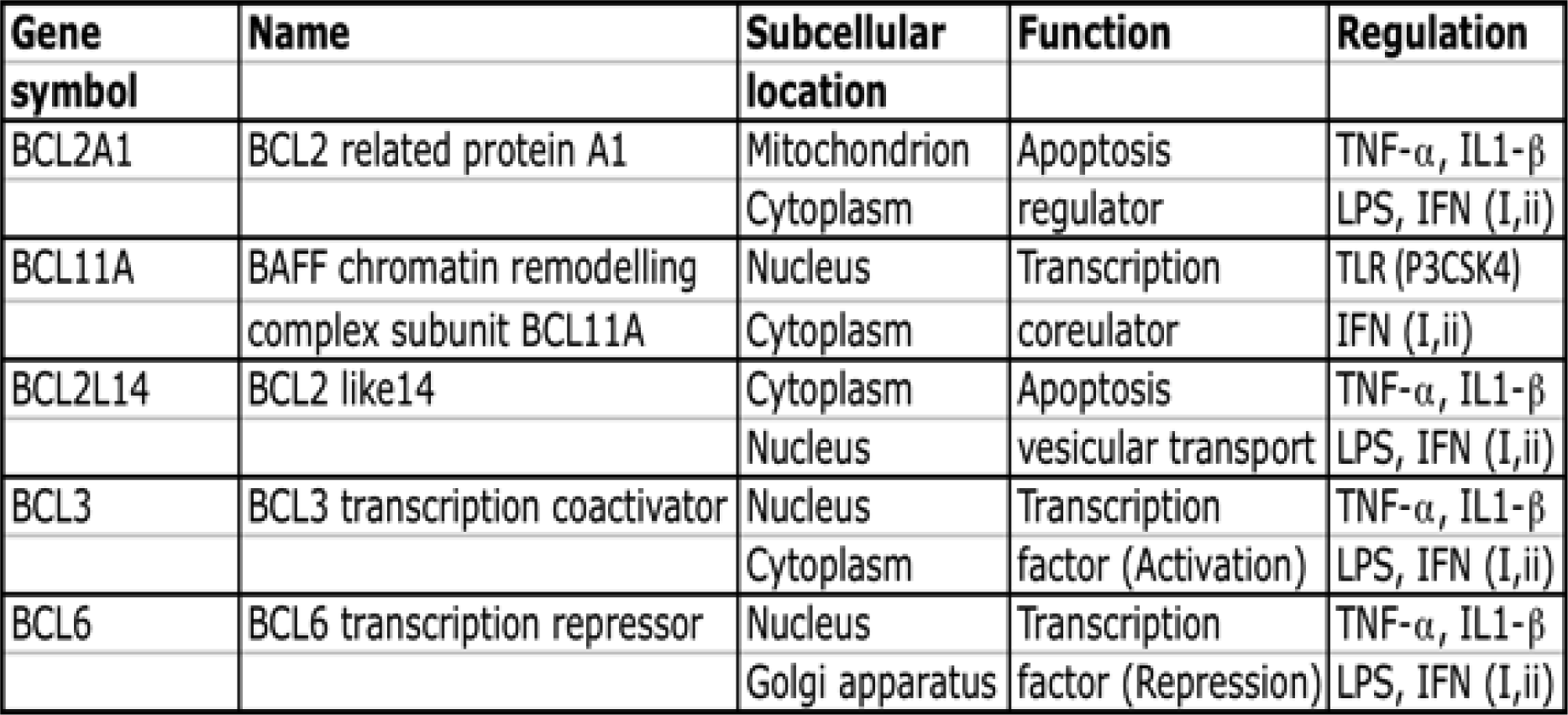
Expression, subcellular location function, and regulation of BCL2 family members.

**Figure 1.**
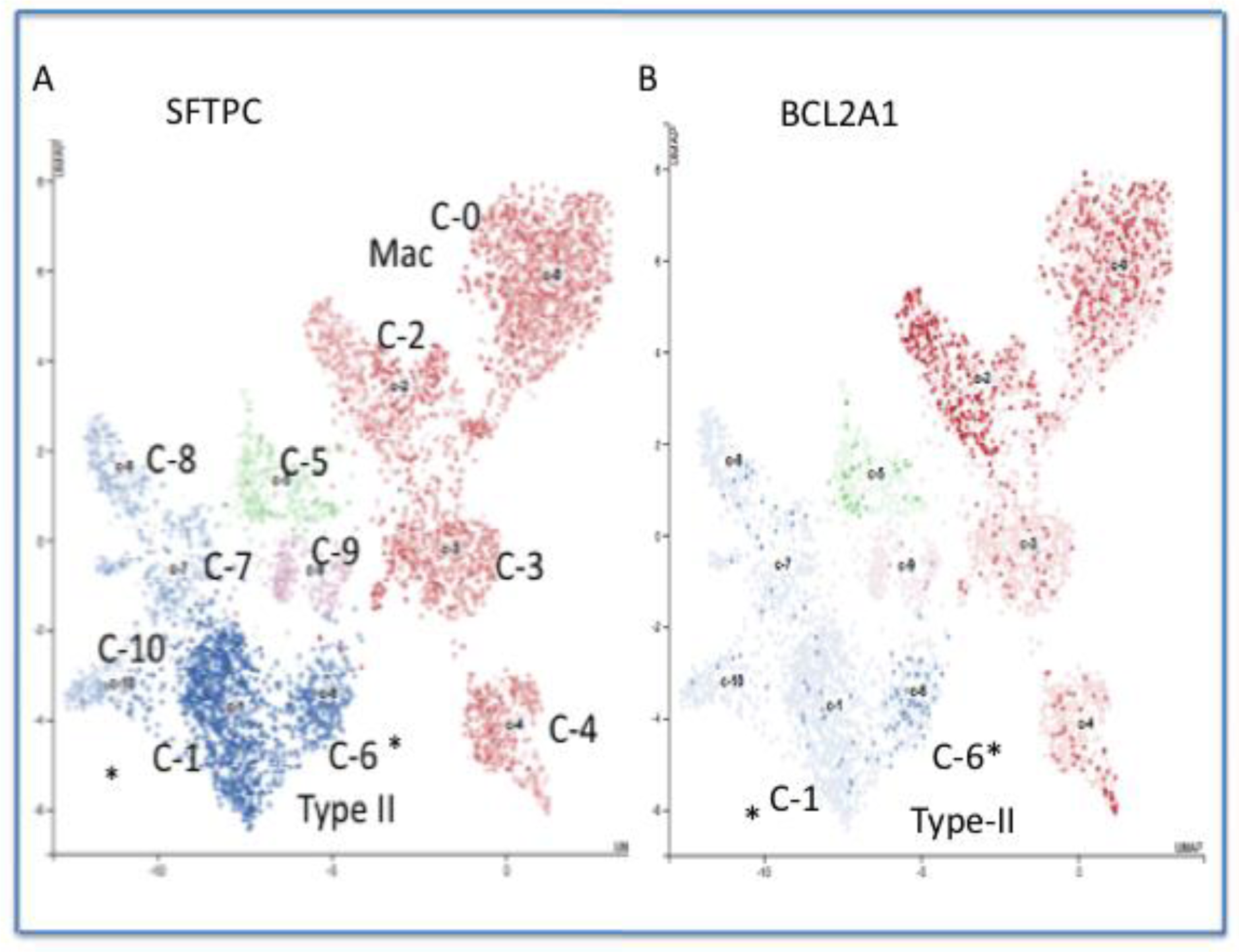
Cell-specific RNA expression of SFTPC (AT2 cell marker) and BCL2A1 in the Human Protein Cell Atlas database. RNA expression profiles were retrieved from the human lung tissue dataset. Each point represents a single cell, colored according to cell type. SFTPC expressed in the C-2 and the C-6 AT2 cell populations while BCL2A1was expressed predominantly in the C-6 AT2 cell population. AT2 (type II) cells were indicated by an asterisk. Mac refers to the C-0 and the C-2 macrophage cell population.

**Figure 2.**
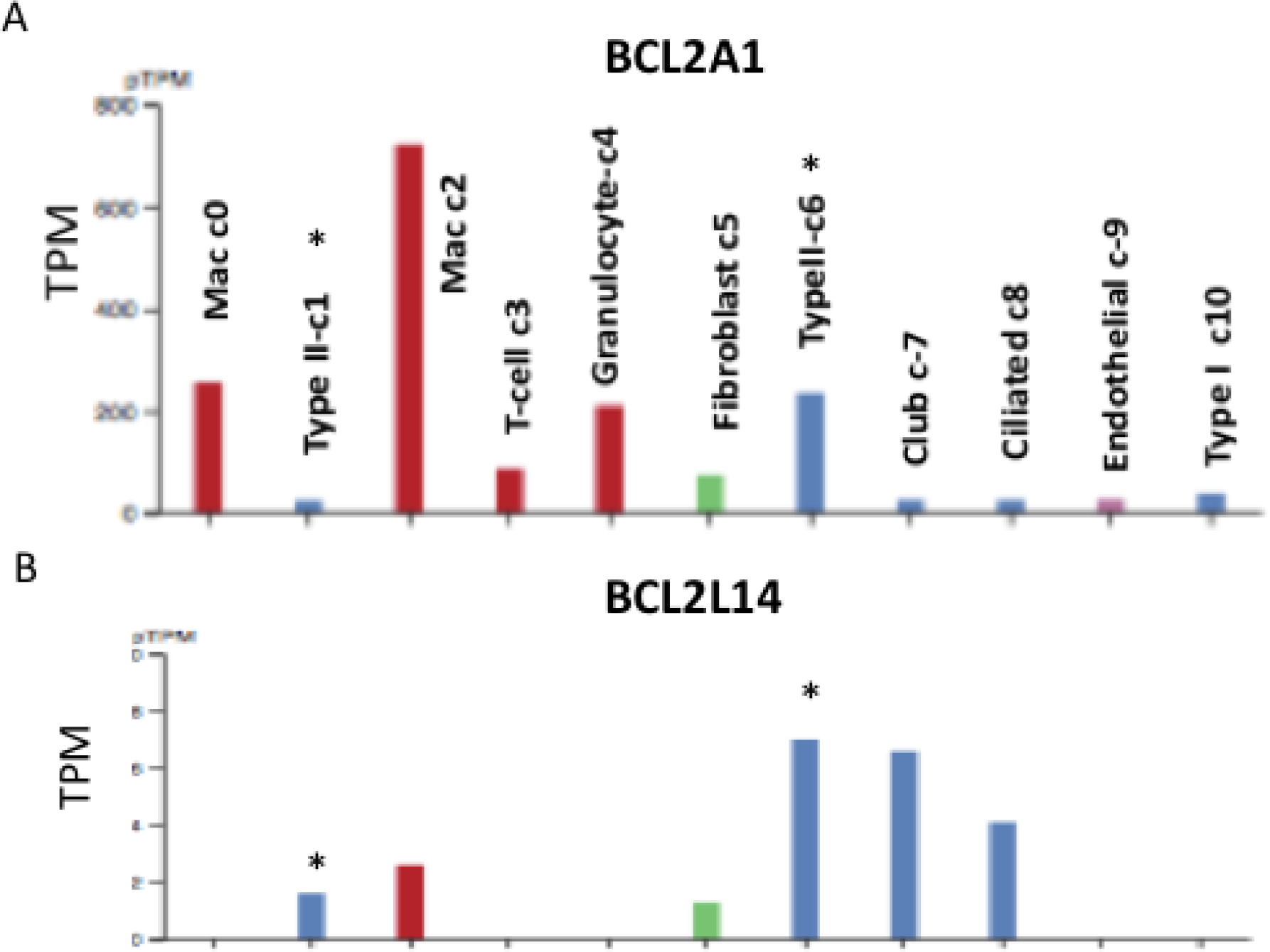
Cell-specific expression of BCL2A1 and BCL2L14 RNA levels in the Human Protein Cell Atlas database. RNA expression profiles were retrieved from the human lung tissue dataset. Human lung C-1 and the C-6 AT2 cell populations were indicated by an asterisk. Transcripts per million reads or TPM were shown.

**Figure 3.**
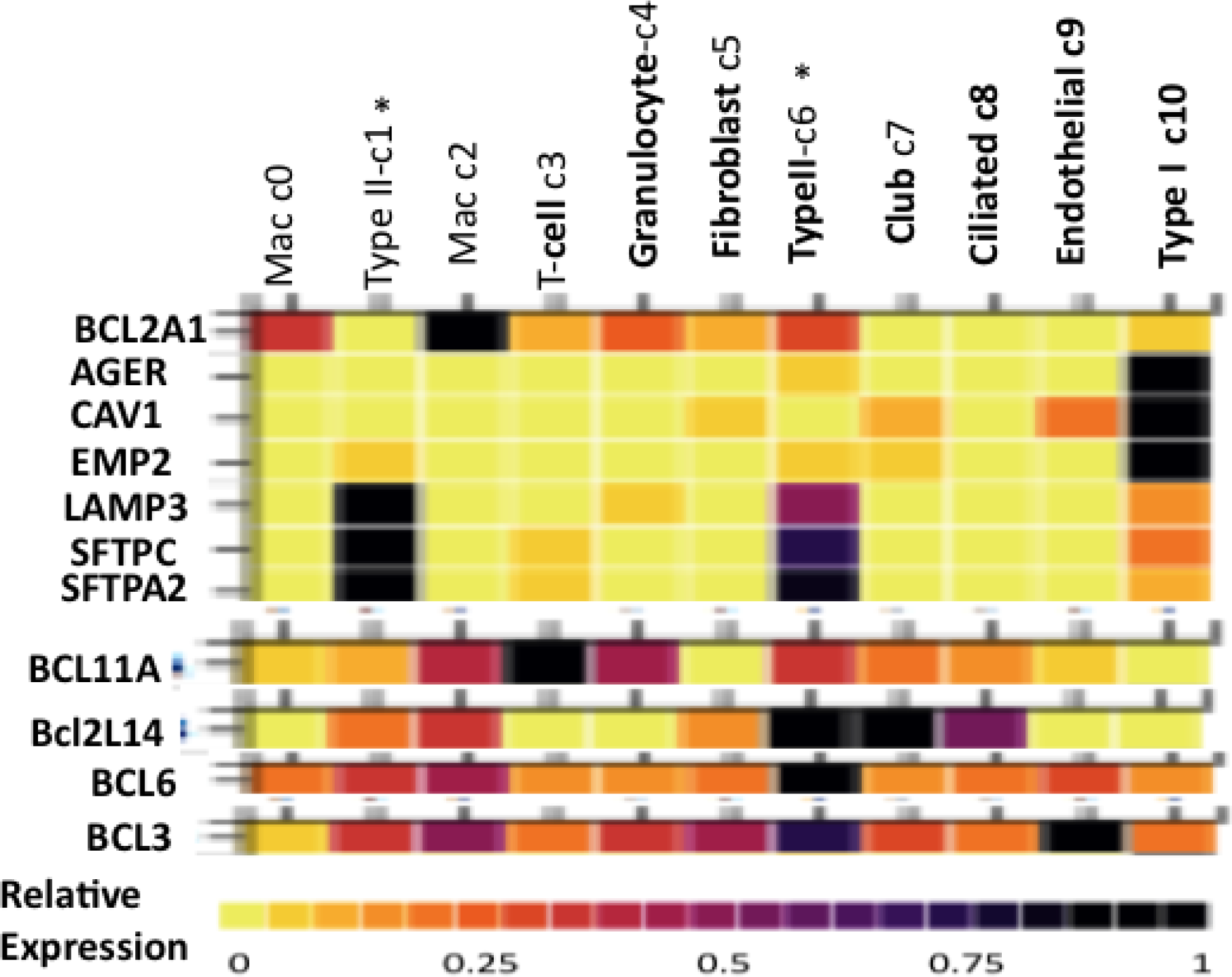
Heatmap representation of the RNA levels of selected members of BCL-2 related family in the distinct cell populations of the human lung. RNA expression profiles were retrieved from the human lung tissue dataset and represented as a heatmap. Human lung type II C-1 and the C-6 cell population were indicated by an asterisk. Expression levels of lung AT2 markers SFTPC, SFTPA2, LAMP3, and AT1 markers AGER, CAV1, and EMP2 were shown for comparison.

### 3.2 Tissue distribution, expression and regulation of BCL2A1

Tissue distribution analysis in a large set of human tissues in the GTEx database (Broad Institute, Cambridge, MA) confirmed that BCL2A1 was highly expressed in the human whole blood, lung. and Epstein-Barr virus (EBV)-transformed B lymphocytes (Figure 4). BCL2A1 was originally identified as an apoptosis regulator in the interleukin-3 (IL-3) dependent murine myeloid cell line (40). Furthermore, BCL2A1 was up-regulated by tumor-promoting phorbol esters (PMA) and inflammatory cytokines TNF- α and IL-1β in human endothelial (HUVEC) and human microvascular (HMEC1) cell lines (41–43). These results suggest BCL2A1 may function in the response of cells to inflammatory signals and maintain endothelial survival during viral infection and inhibit cell death during growth factor deprivation. Transcriptional regulation of BCL2A1 by nuclear factor kappaB (NF-KB) in response to TNF-α and IL-1β was reported (44). The intracellular location of BCL2A1 includes the outer membrane of the mitochondria and cytoplasm (Table 1). BCL2A1 blocks caspase activation and release of cytochrome c from mitochondria and inhibits intrinsic apoptosis pathway (45). Annotation in the TCGA cancer database revealed that BCL2A1 was transcriptionally misregulated in cancers like acute myeloid leukemia (data not shown). Interrogation of multiple microarray datasets revealed that BCL2A1 mRNA levels were increased 4-6 fold by inflammatory cytokines (TNF-α, ILl-β), TLR ligands (LPS, P3CSK4) and Vitamin D3. In contrast, transforming growth factor (TGFB1) and steroid hormone Dexamethasone (Dex) suppressed BCL2A1 mRNA levels (Figure 5A). Immgen microarray data revealed high basal expression of Bcl2a1 in mouse alveolar macrophages and poly-IC (a double-stranded RNA mimetic and inducer of type I interferon) further enhanced expression by 2-fold at RNA levels (Figure 5B). These results prompted a detailed study on the regulation of BCL2A1 by respiratory viruses and type I interferon signaling in the human lung epithelial cells.

**Figure 4.**
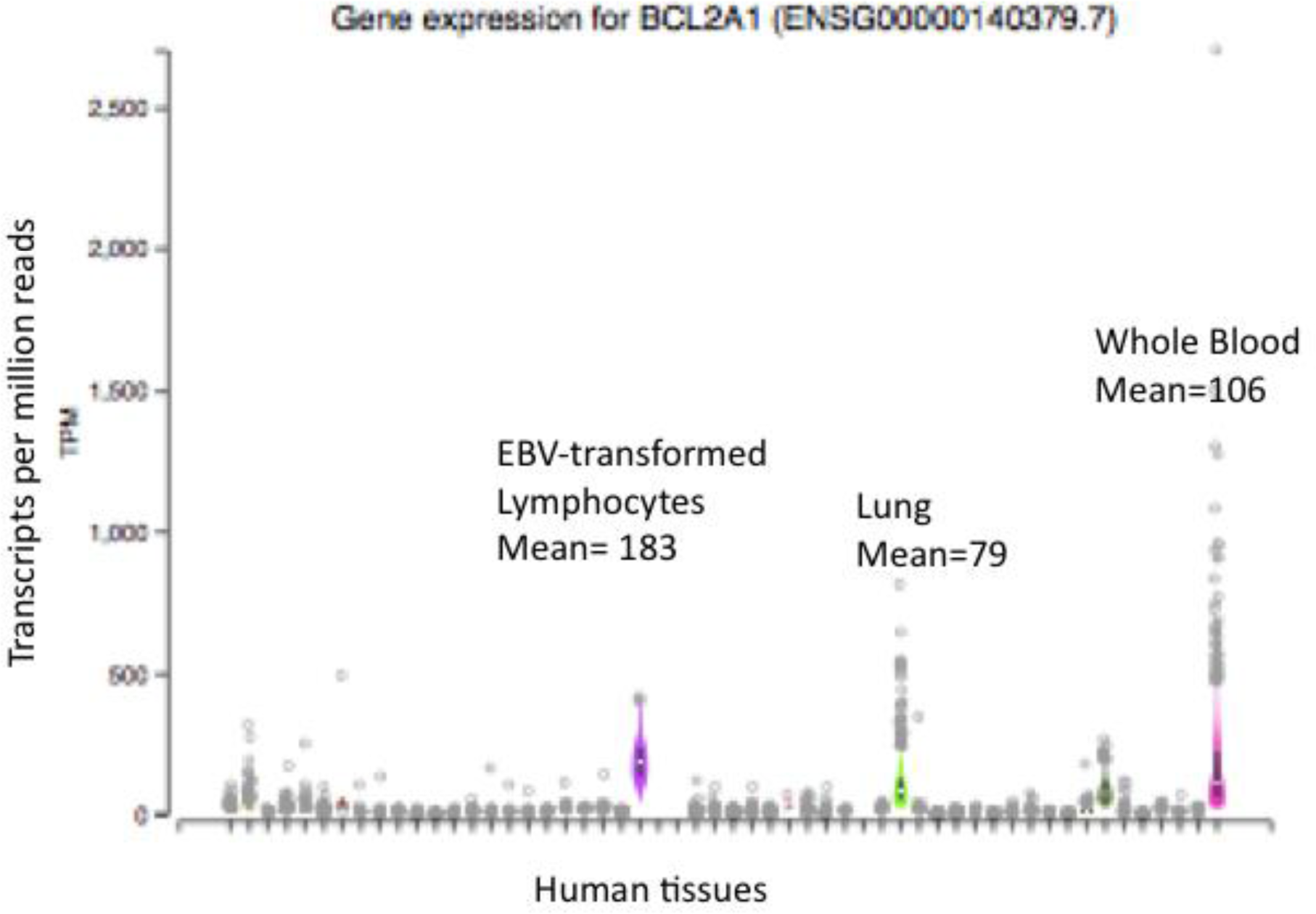
Expression levels of BCL2A1 RNA in human tissues in the GTEx database. High levels of BCL2A1 expression in the whole blood, lungs, and Epstein-Barr virus (EBV)-transformed lymphocytes were shown.

**Figure 5.**
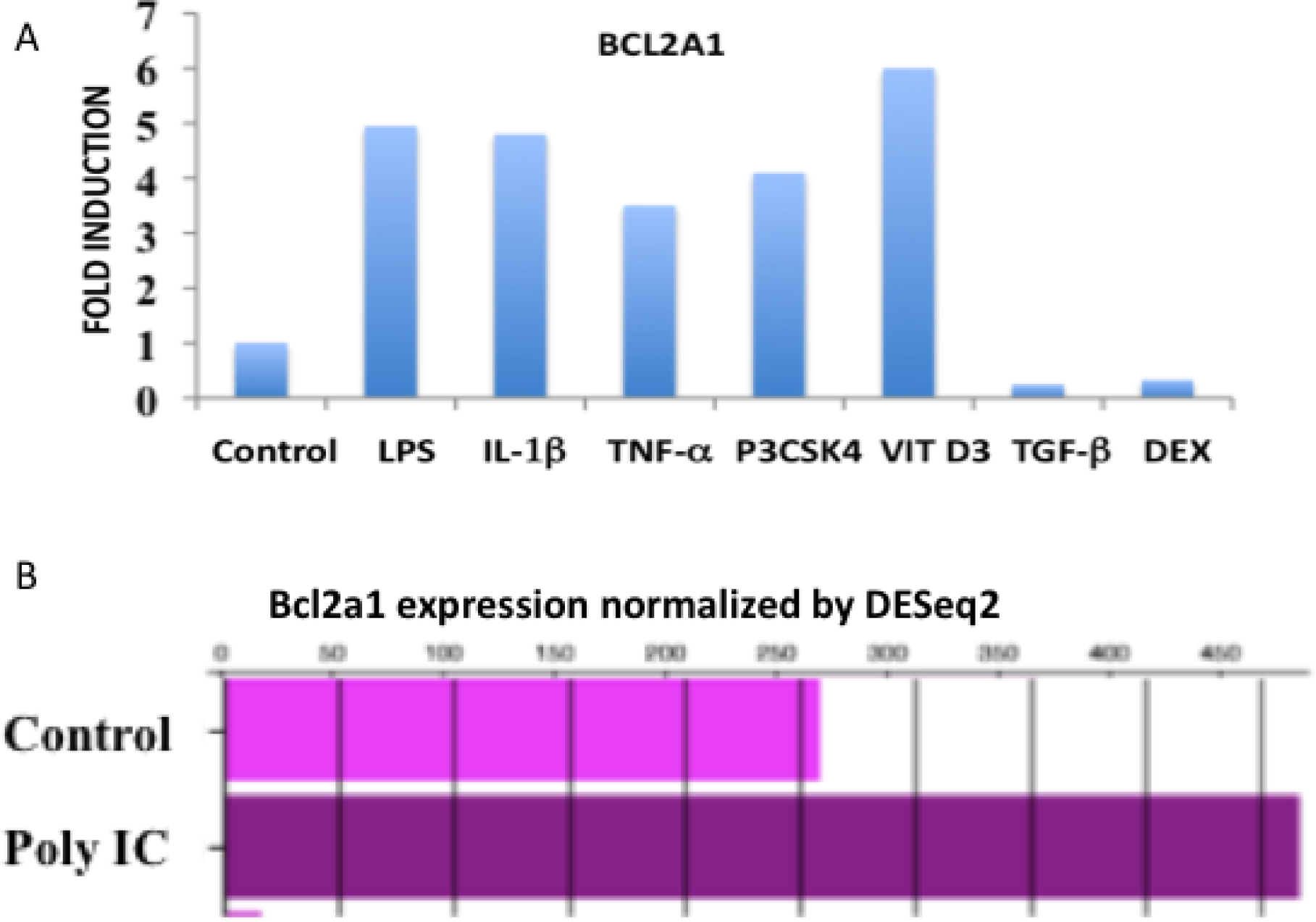
Regulation of BCL2A1 mRNA levels by cytokines, TLR ligands, and steroid hormones. (A) Relative levels of BCL2A1 mRNA in liver cells after treatment with cytokines (IL-1β, TNF-α, or TGF-β), TLR ligands (LPS, or P3CSK4) or steroid hormones (Vitamin D3, or DEX). (B) Relative levels of Bcl2a1 mRNA in the control alveolar macrophages of C57/BL6 mice and after treatment with type I interferon inducer (poly-IC).

### 3.3 Regulation of BCL2A1 expression by coronaviruses

Coronaviruses such as SARS CoV-2 require human lung epithelial cell membrane proteins ACE2 and TMPRSS2 for productive viral infection and facilitate viral entry and release of the viral RNA into the host cell cytoplasm (2, 5). Recognition of viral RNA by the host cell Toll-like receptor 3 (TLR3) pathway results in the phosphorylation and activation of IRF3 and activation of type I interferon gene transcription, protein synthesis, and secretion. IFN α/β binds to its receptor on the surface of infected or neighboring cells in an autocrine or paracrine fashion to activate the Jak-Stat pathway to induce interferon-stimulated genes and establish the antiviral state (11, 12). This innate immune response is the body’s first line of defense against the recognition and clearance of respiratory viruses (1,2,5). Transcriptome analysis in widely used human lung epithelial cell lines such as Calu-3, A549, and NHBE1 in response to coronaviruses was deposited in Pubmed/NCBI (Geo datasets) and Immgen Covid 19 skyline database. Interrogation of Microarray datasets revealed that the mRNA levels of BCL2A1 and other members of the BCL-2 family such as BCL2L14, BCL3, and BCL6 were induced by SARS CoV-2 infection but not by mock-infection in Calu-3 cells. This induction was observed within 24 hours after infection (Figure 6A). Furthermore, interferon-beta 1 (IFNB1) and transcription factors STAT1, STAT2, and IRF7 were also induced at mRNA levels by SARS CoV-2 infection in Calu-3 cells (Figure 6B). These results suggest that the production of IFN beta and autocrine activation of the Jak-Stat pathway plays a major role in the induction of interferon-stimulated genes. In contrast, SARS-CoV-2 infection of A549 failed to induce IFNB1 and interferon-stimulated gen expression (Figure 7A). Previous studies have shown that ACE2 functions as the virus entry receptor and facilitates the entry of SARS-CoV-2 into lung epithelial cells (46). Expression of ACE2 in A549 cells rescued SARS-CoV-2 -mediated induction of IFNB1, STAT1, STAT2, and IRF9 mRNA (Figure 7A). Furthermore, ACE2R expression dramatically increased the expression of BCL2A1, BCL3, and BCL6 mRNA after SARS-CoV-2 infection in A549/ACE2 cells (Figure 7B). These results suggested that BCL2A1 and the expression of selected BCL-2 family members was dependent on IFNB1 production by SARS-CoV-2 in A549 cells and required the expression of the virus entry receptor ACE2. In contrast, time-course analysis of BCL2A1 and BCL2L14 RNA in HAE human lung epithelial cells revealed maximal induction at 72 hours after SARS-CoV-1 infection (Table 2 and Table 3). However, these microarray datasets were generated in different cell lines and under different experimental conditions and may not be directly comparable to each other.

**Figure 6.**
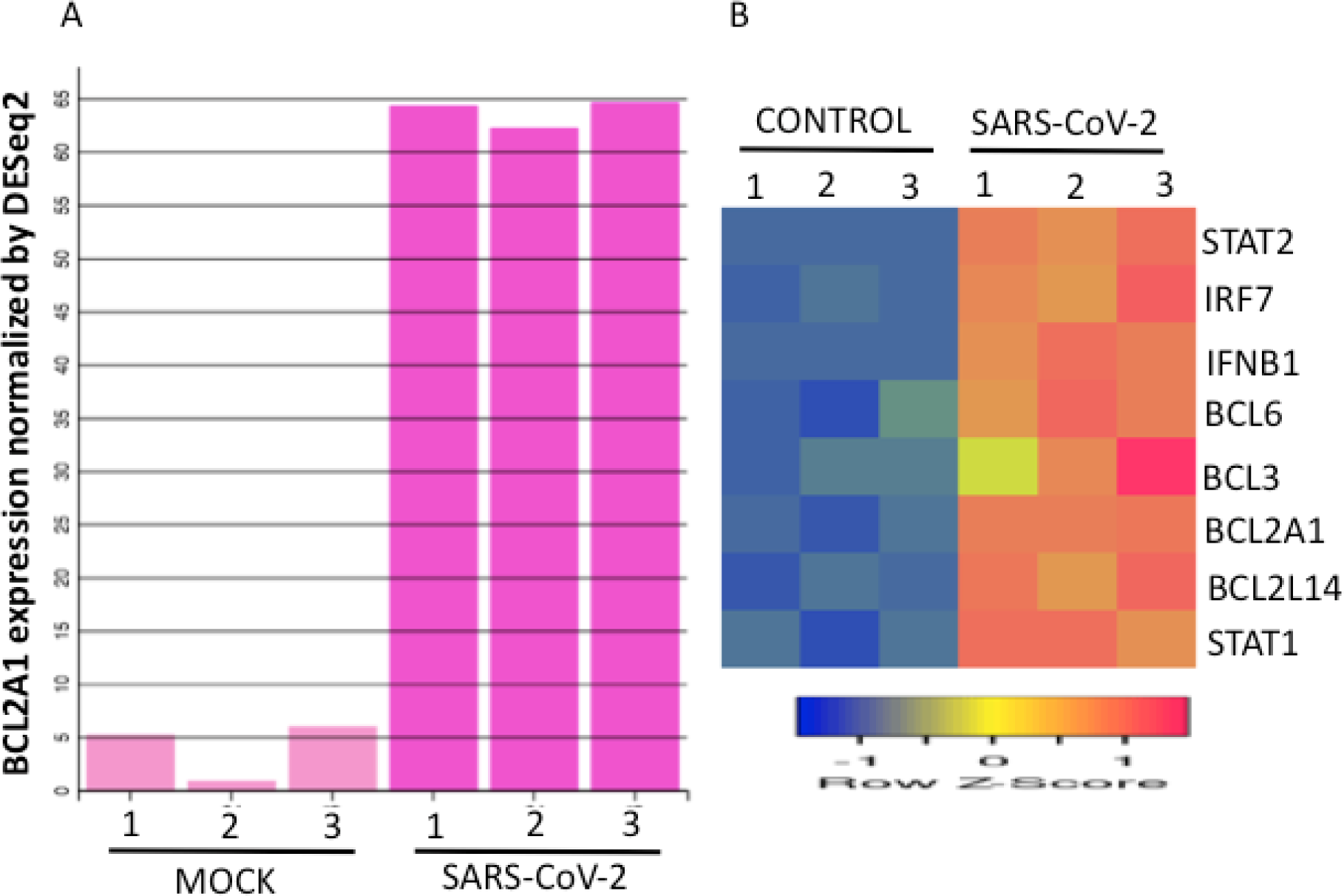
Regulation of the BCL-2 related family member RNA levels by SARS CoV-2 in Calu-3 cells. (A) Calu-3 cells were mock-infected or infected with the SARS-CoV-2 for 24 hours. BCL2A1 RNA expression levels were normalized by the DEseq2 method (B) Regulation of interferon-stimulated genes and BCL-2 related family members by the SARS CoV-2 in Calu-3 cells. Mock-infection for 24 hours was used as control. Data from three samples for each condition were shown. RNA expression levels were normalized by the DEseq2 method. Sample IDs 1, 2, and 3 from the Immgen dataset were represented in the figure.

**Figure 7.**
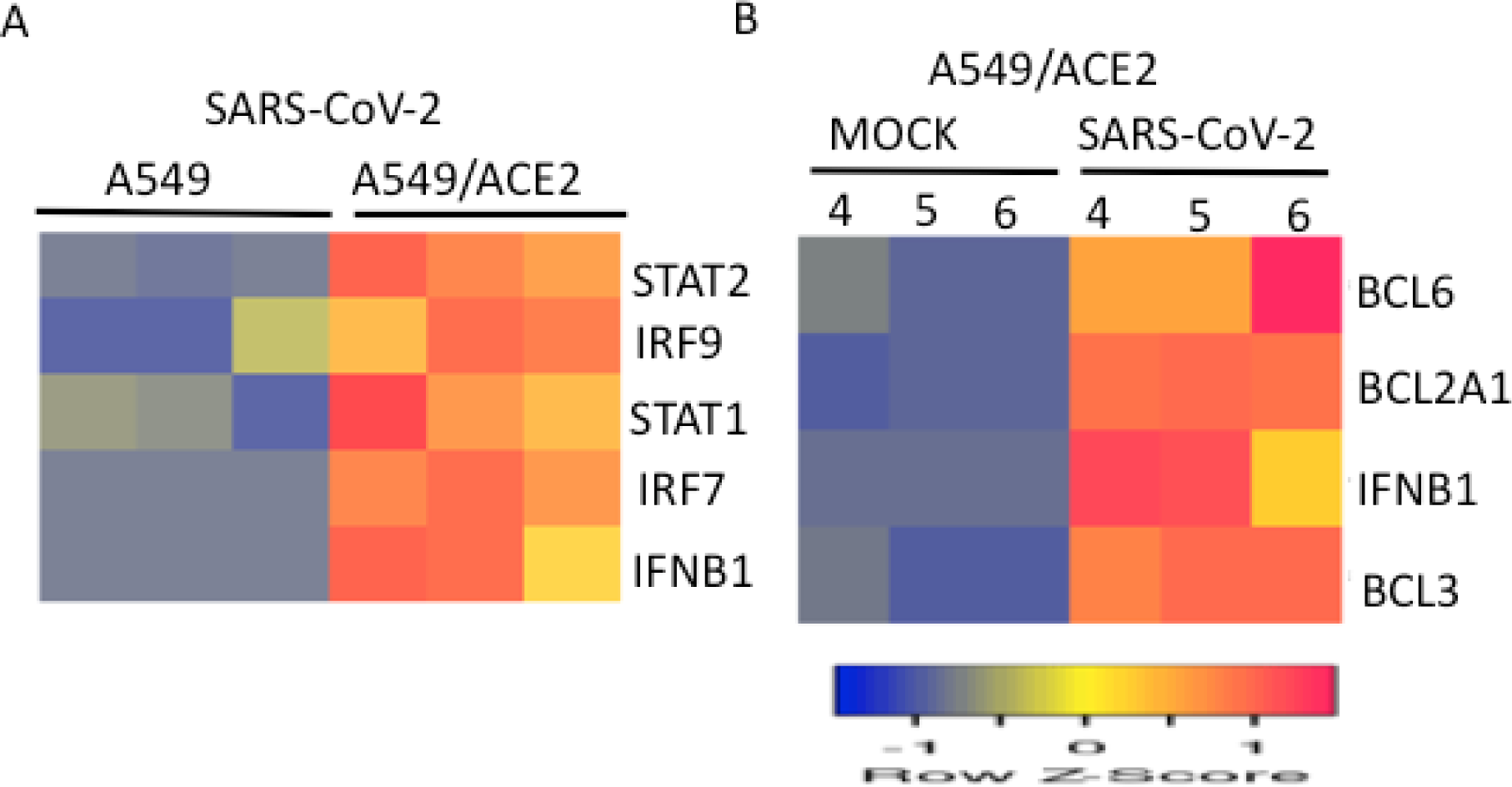
Regulation of the interferon-stimulated genes and members of BCL2-related family RNA levels by SARS-CoV-2 in A549 or A549 cells expressing ACE2 receptors. (A) A549 or A549 cells expressing ACE2 receptors were mock-infected or infected with the SARS-CoV-2 virus for 24 hours and RNA fold-induction of IFN-regulated transcription factors from three samples for each condition was shown. (B) The expression levels of BCL2A1, BCL3, BCL6, and IFNB1 RNA in A549 cells expressing ACE2 receptors after mock or virus infection were normalized by the DEseq2 method. Data from three samples for each condition were shown. Sample IDs 4, 5, and 6 from the Immgen dataset were represented in the figure.

**Table 2.**
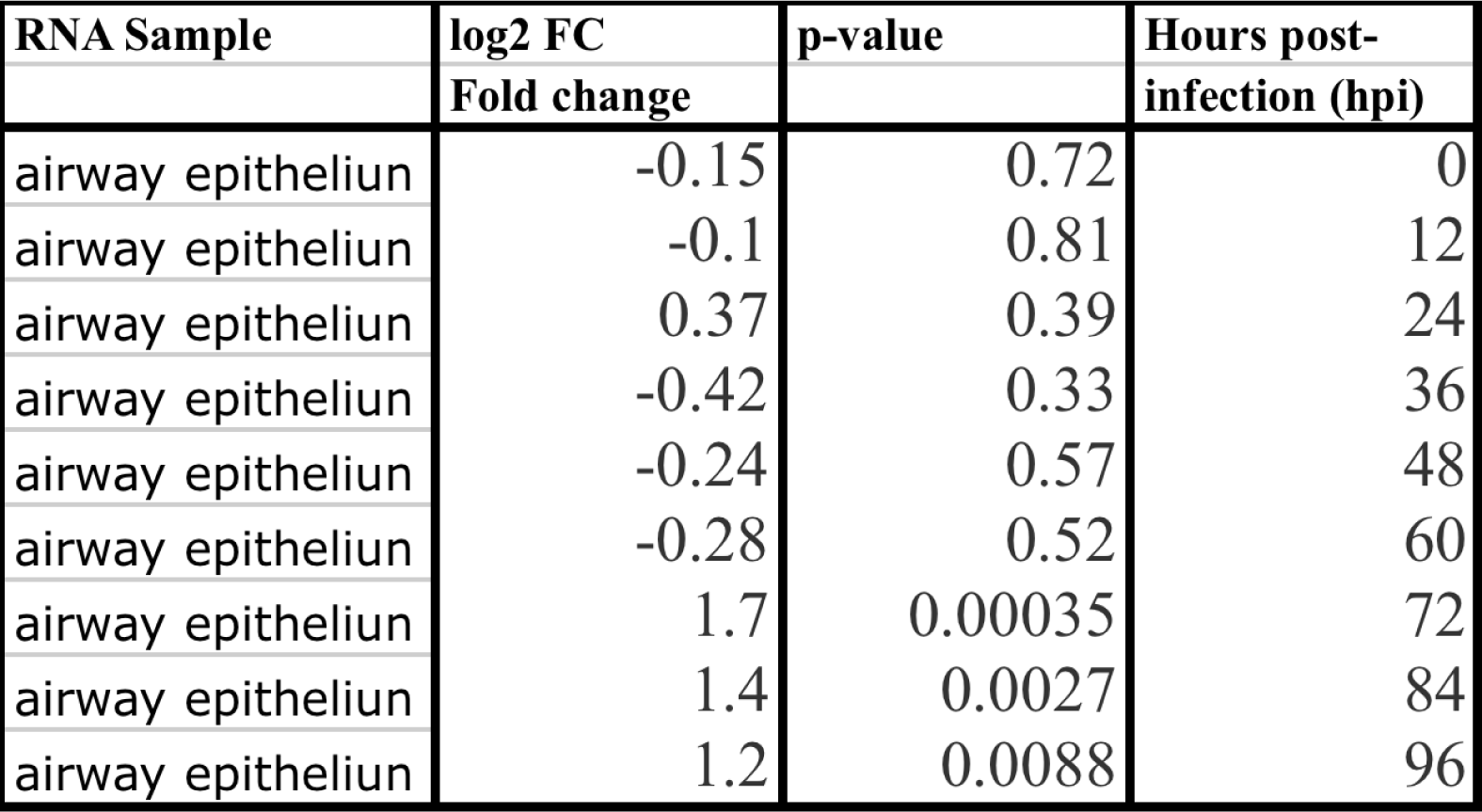
BCL2A1 regulation by SARS-CoV-1 infection in HAE human lung epithelial cells.

| RNA Sample | log2 FC | p-value | Hours post-infection (hpi) |
| --- | --- | --- | --- |
|  | Fold change |  |  |
| airway epitheliun | -0.15 | 0.72 | 0 |
| airway epitheliun | -0.1 | 0.81 | 12 |
| airway epitheliun | 0.37 | 0.39 | 24 |
| airway epitheliun | -0.42 | 0.33 | 36 |
| airway epitheliun | -0.24 | 0.57 | 48 |
| airway epitheliun | -0.28 | 0.52 | 60 |
| airway epitheliun | 1.7 | 0.00035 | 72 |
| airway epitheliun | 1.4 | 0.0027 | 84 |
| airway epitheliun | 1.2 | 0.0088 | 96 |

**Table 3.** BCL2L14 regulation by SARS-CoV-1 infection in HAE human lung epithelial cells.

| **RNA Sample** | **log2 FC** | **p-value** | **Hours post-infection (hpi)** |
| --- | --- | --- | --- |
|  | **Fold change** |  |  |
| airway epitheliun | -0.23 | 0.3 | 0 |
| airway epitheliun | 0.29 | 0.2 | 12 |
| airway epitheliun | 0.12 | 0.58 | 24 |
| airway epitheliun | -0.14 | 0.54 | 36 |
| airway epitheliun | 0.27 | 0.23 | 48 |
| airway epitheliun | -0.13 | 0.57 | 60 |
| airway epitheliun | 1.3 | 1.40E-06 | 72 |
| airway epitheliun | 1.1 | 9.90E-06 | 84 |
| airway epitheliun | 0.89 | 0.00028 | 96 |

### 3.4 Regulation of BCL2A1 expression by influenza and other respiratory viruses

Respiratory virus infection activates the production of antiviral cytokines like interferons as well as pro-inflammatory cytokines like TNF-α, IL-1β and IL-6 by lung resident immune cells and alveolar epithelial cells (47). The influenza A virus A (IAV) belongs to the Orthomyxoviridae family of viruses and is responsible for seasonal epidemics and more infrequently global pandemics (48). Continuous mutation, reassortment in intermediate hosts, and antigenic drift contribute to the variability of different strains and are responsible for the severity and pathogenicity of the virus (49). In addition to the structural and replication machinery, the virus encodes a non-structural protein (NS1) that is a potent inhibitor of IRF3 activation, type I interferon production, and Stat1 activation resulting in attenuation of interferon-stimulated gene expression (50, 51). Induction of BCL2A1 expression in NHBE bronchial epithelial cells was dependent on NS1 in the virus. Wild-type IAV failed to induce BCL2A1 expression in NHBE cells. In contrast, virus infection with a deleted NS1 resulted in the induction of BCL2A1 expression (Figure 8A). Respiratory syncytial virus (RSV) infection occurs in children while human parainfluenza virus (HPIV3) infections are common in children, immunocompromised adults, and the elderly. RSV and HPIV3 viruses induced BCL2A1 expression in A549 cells (Figure 8B). These studies suggest that multiple factors such as host-encoded factors (ACE2, TMPRSS2), virus-encoded factors (NS1) and cell type play an important role in determining viral pathogenesis. A snapshot summary of BCL2A1 regulation by influenza and coronaviruses was presented with red ovals representing induction and blue ovals representing suppression in separate microarray experiments (Figure 9).

**Figure 8.**
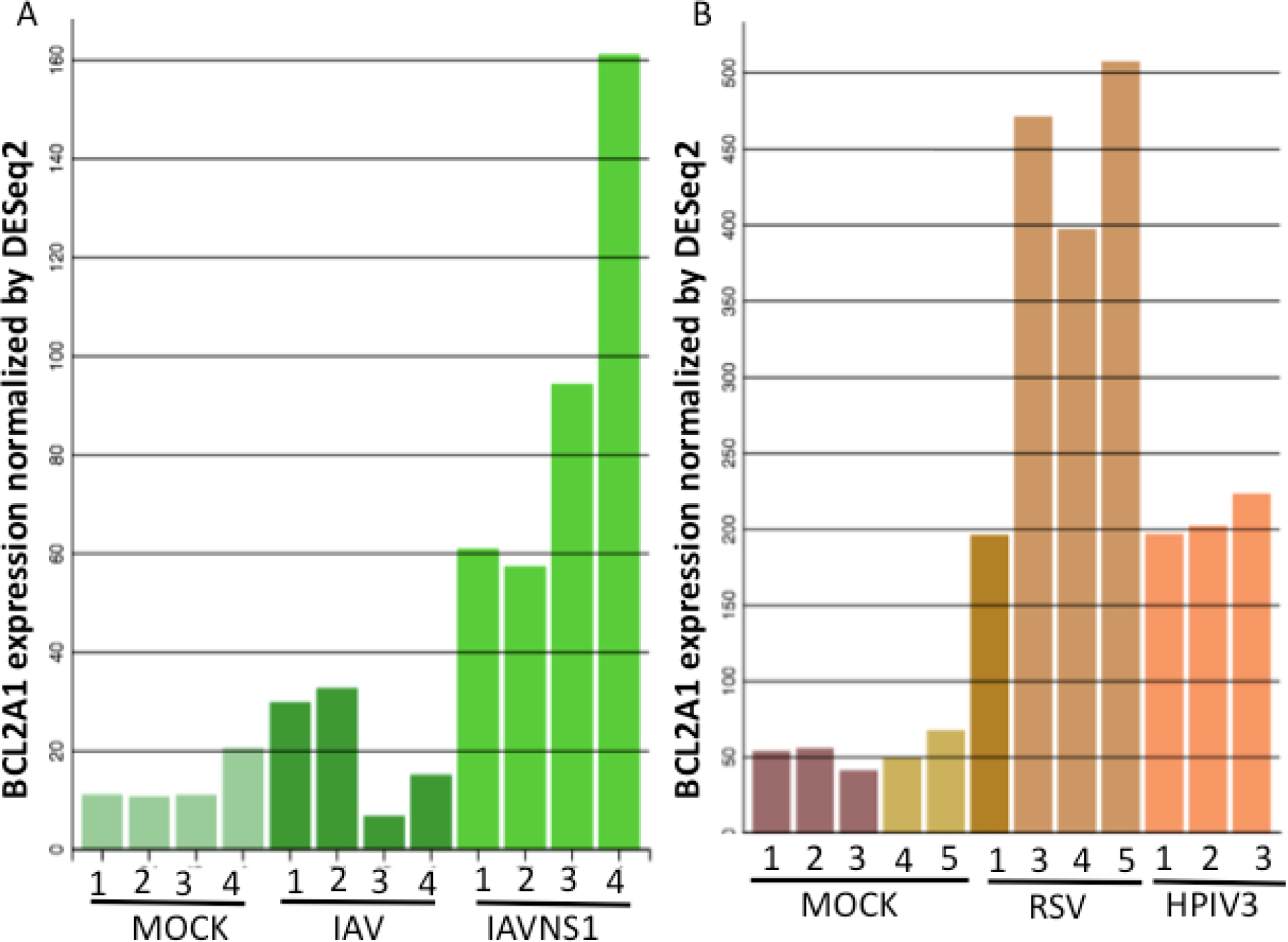
Regulation of BCL2A1 mRNA levels by the influenza viruses (IAV), respiratory syncytial virus (RSV), and the human parainfluenza virus (HPIV3) (A) Human NHBE1 bronchial epithelial cells were mock-infected or infected with the influenza virus (IAV) or influenza virus lacking the NS1 (IAVNS1) for 12 hours. Results from four samples (IDs 1, 2, 3, and 4 from the Immgen dataset) for each condition were shown (B)) Human A549 lung cells were mock-infected or infected with the RSV or HPIV3 for 24 hrs. BCL2A1 mRNA levels were normalized by the DESeq2 method. Data from 3-5 samples for each condition was shown. Sample IDs 1, 2, 3, 4, and 5 from the Immgen dataset were represented in the figure.

**Figure 9.**
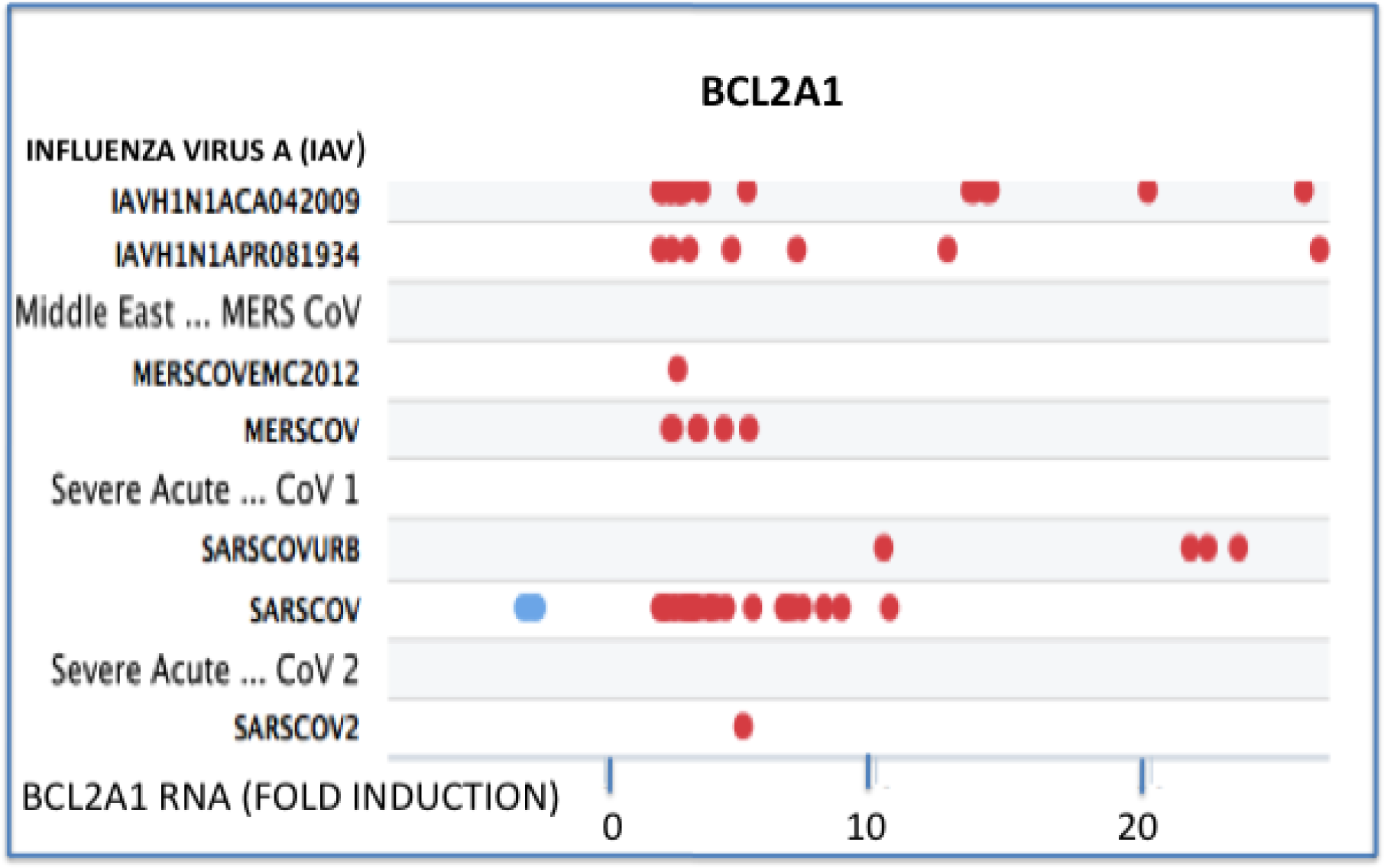
A snapshot of the regulation of BCL2A1 mRNA levels by respiratory viruses as represented in the microarray datasets. Induction or suppression of BCL2A1 mRNA levels by respiratory viruses (IAV, MERS, SARS CoV-1, and the SARS CoV-2) was represented by red and blue ovals, respectively. Each oval represents a separate microarray experiment.

### 3.5 Regulation of BCL2A1 expression by type I Interferon signaling

There are two distinct modes of type I Interferon (IFN-α/β) signaling- a rapid phosphorylation switch mediated by the Jak-Stat cascade and a graded output of transcription factor induction at mRNA level that enforces temporal regulation (52). The first transcriptional switch takes place within 30 minutes and drives primary interferon-stimulated gene expression mediated by Stat, Stat2, and IRF-9 (ISGF2) transcription factor complexes or Stat homo or heterodimers (53). The second transcriptional switch takes place several hours later and is mediated by interferon-inducible transcription factors that sustain secondary and tertiary rounds of interferon-stimulated gene expression (52). Interrogation of microarray datasets revealed that IFN-β treatment induced STAT1 and STAT2 transcription factor mRNA as well as BCL2A1 and BCL2L14 RNA expression in 8 hours in A549 human lung epithelial cells (Figure 10A).

**Figure 10.**
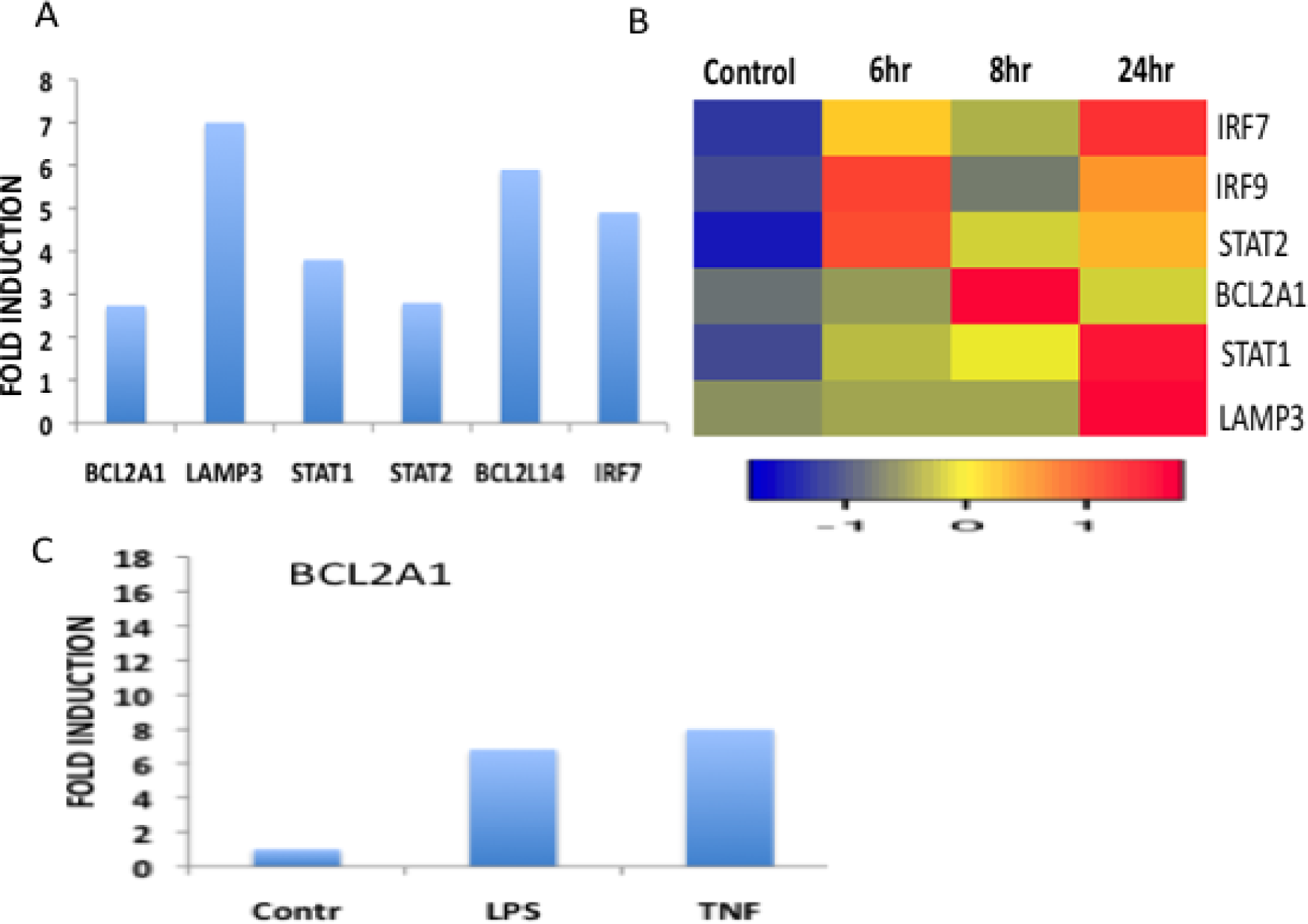
Regulation of the RNA levels of interferon-stimulated genes (ISG) and BCL-2 family members by interferon-beta (IFN-β) treatment in A549 human lung epithelial cells (A) Regulation of RNA levels of ISG by IFN-β treatment for 8 hours in A549 cells (B) Regulation of RNA levels of ISG by IFN-β treatment for 6, 8, or 24 hours in A549 cells (C) Regulation of BCL2 A1 mRNA levels by LPS and TNF-α in A549 cells. Gene expression data were retrieved from microarray datasets.

Times-course microarray data revealed that BCL2 A1 expression was induced transiently at 8 hr and decreased to basal levels by 24 hours (Figure 10B). In contrast, BCL2A1 was rapidly induced by inflammatory stimuli TNF-α, IL-1β, LPS within 6 hours in A549 cells (54; Figure 10 C). These studies suggest that BCL2 A1 RNA was induced in a distinct temporal fashion in response to inflammatory signals and type I interferon in A549 cells. In contrast, in NHBE bronchial epithelial cells IFN-β rapidly and transiently induced BCL2A1 mRNA levels within 4-6 hours and the levels declined by 12 hours (Figure 11A). Transcriptional regulation by type 1 interferons is mediated by STAT1, STAT2, and IRF9 binding to regulatory elements known as interferon-stimulated response element (ISRE) or STAT1 and STAT3 homo-or heterodimers binding to another regulatory element known as gamma-activated sequence (GAS) located near transcription start site (TSS) of interferon-stimulated genes (52, 53). Transcription factor binding site data in the promoter region of BCL2A1 in the Interferome database revealed that there were no ISRE or GAS elements within the 1.5 Kb upstream of the transcription start site. Interestingly, NF-kB, IRF1, and IRF7 binding sites were detected (Figure 11B). Chromatin immunoprecipitation experiments revealed that Stat1 bound to the proximal promoter region of the mouse Bcl2a1 in response to interferon-γ. However, the exact location of the sequence element or Stat1 binding site in the promoter remains to be determined (55). Immgen microarray data revealed that IFN-α treatment for 2 hours had no significant effect on Bcl2a1 mRNA levels in B lymphocytes, dendritic cells, natural killer cells and NKT cells (Figure 12A). In contrast, in granulocytes and macrophages, IFN-α has no effect while IFN-γ significantly suppressed Bcl2a1 at mRNA levels (Figure 12B). Interestingly, granulocyte macrophage-colony stimulating factor (GM-CSF) stimulated the Bcl2a1 expression while co-treatment of GM-CSF with IFN-γ suppressed the Bcl2a1 expression in mouse CD11b-Grl^+ (high)^ myeloid-derived suppressor cells (55).

**Figure 11.**
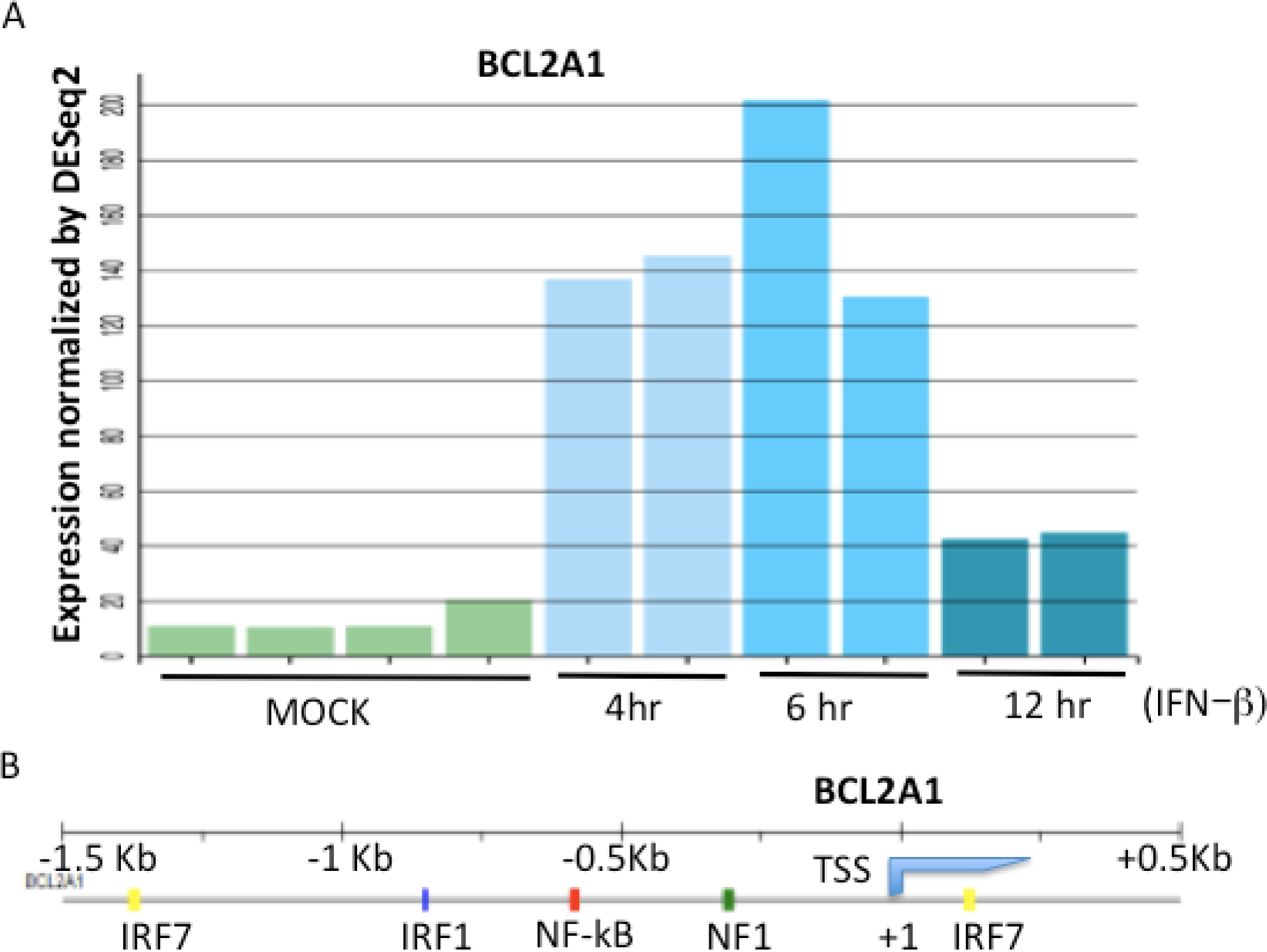
Regulation of BCL2A1 mRNA levels by interferon-beta (IFN-β) treatment in human NHBE lung epithelial cells. (A) Regulation of BCL2A1 RNA levels by IFN-β treatment for 4, 6, or 12 hours in human NHBE cells (B) Schematic representation of the human BCL2A1 gene promoter. The approximate location of the NF-kB, IRF7, IRF1 and NF1 binding sites was shown. TSS represents the transcription start site.

**Figure 12.**
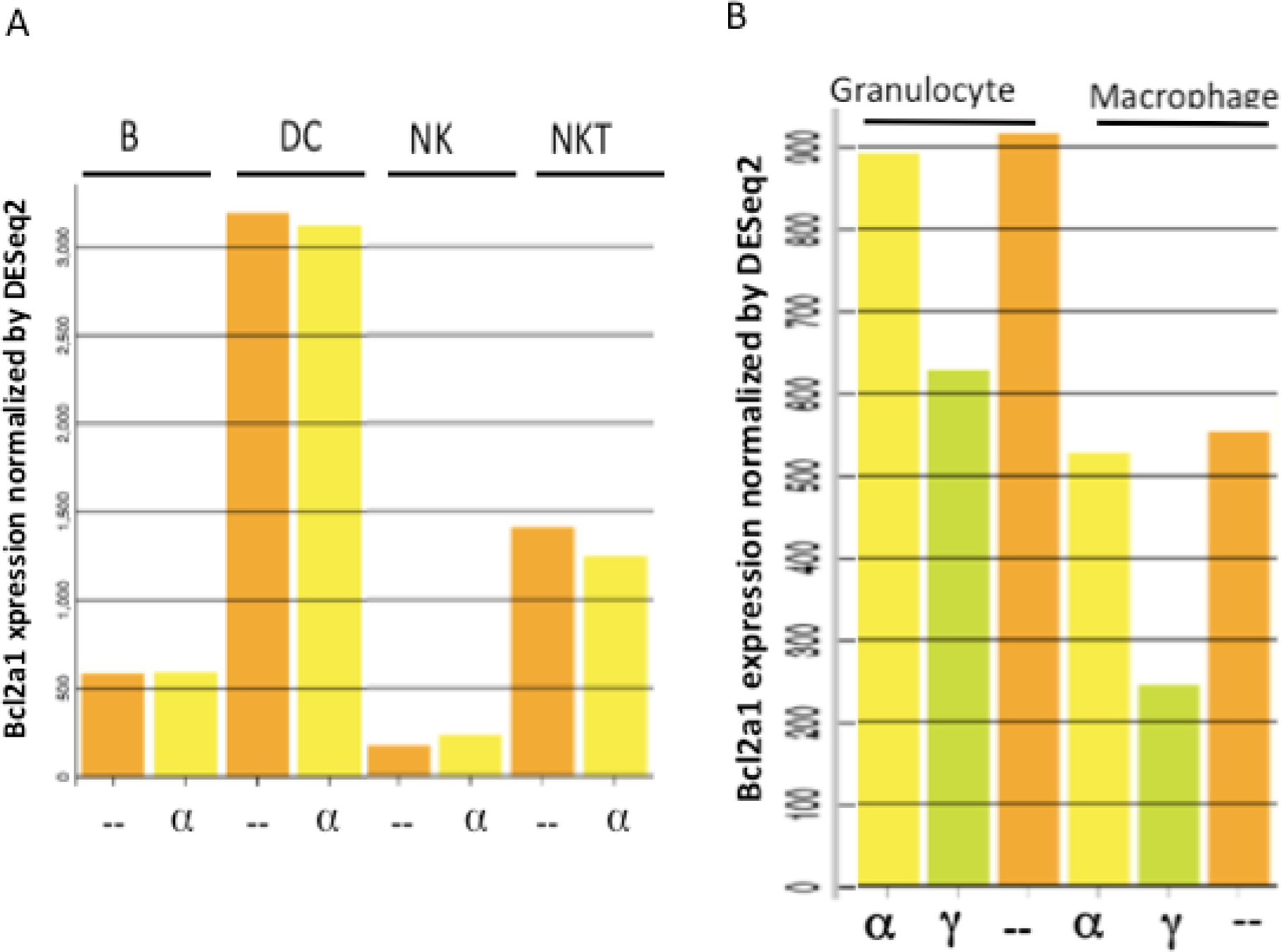
Differential regulation of BCL2 A1 RNA levels by interferon-alpha (IFN-α) and interferon-gamma (IFN-γ) in the mouse immune cells. Immgen microarray data of B-lymphocyte (B), dendritic cells (DC), natural killer cells (NK), natural killer T cells (NKT), granulocytes, and macrophage cells that were left untreated, treated with IFN-α or IFN−γ for 2 hours.

### 3.6 Regulation of BCL2 L14 by interferons and respiratory viruses

BCL-G or BCL2L14 is a member of the BCL2 family and highly expressed in the testis and gastrointestinal tract including the colon, small intestine and stomach (56). Loss of BCL2L14 promotes inflammatory bowel disease and inflammation-associated colon cancer (57). TNF-α and IFN-γ synergistically upregulated BCL2L14 and induced apoptosis in colon epithelial cell lines. However, Th1 cytokine-mediated colon epithelial apoptosis was independent of BCL2L14 expression and STAT1- dependent (58). Furthermore, studies in knockout mice and cells suggested that contrary to the earlier claims, BCL-G/BCL2L14 was not involved in pro-apoptotic function (59). BCL2L14 was among several interferon-stimulated genes associated with type I interferon-induced HIV restriction in humans (60). Interestingly, BCL2L14 mRNA levels were also significantly induced by respiratory viruses including influenza A (IAV), coronaviruses SARS-CoV-1 and SARS-CoV-2 (Figure 13A). Interrogation of the interferon microarray data suggested that BCL2L14 was induced by the type I and type II interferons (Figure 13B). IFN-α treatment induced BCL2L14 mRNA levels in A549 lung epithelial cells (61). Proteins interacting with BCLG/BCL2L14 were identified by yeast 2-hybrid analysis as well as mass-spectroscopy and include components of the Transport particle protein (TRAPP) complex involved in vesicular transport in the early secretory pathway between the endoplasmic reticulum (ER) and Golgi apparatus (62, 63). Previous studies have shown that lysosomal–associated membrane protein 3 (LAMP3) involved in vesicular transport was regulated by respiratory viruses and type I interferon signaling (27). These results revealed that vesicular transport in lung epithelial cells may be a common target of innate immunity regulation by type I interferon signaling and respiratory viruses. Transcriptional regulation analysis in TRANSFAC database suggested that regulatory elements in the BCL2L14 gene promoter included binding sites for STAT1, IRF3 and IRF8 (Figure 13C). In contrast to BCL2A1, dexamethasone treatment up-regulated BCL2L14 expression (supplementary data).

**Figure 13.**
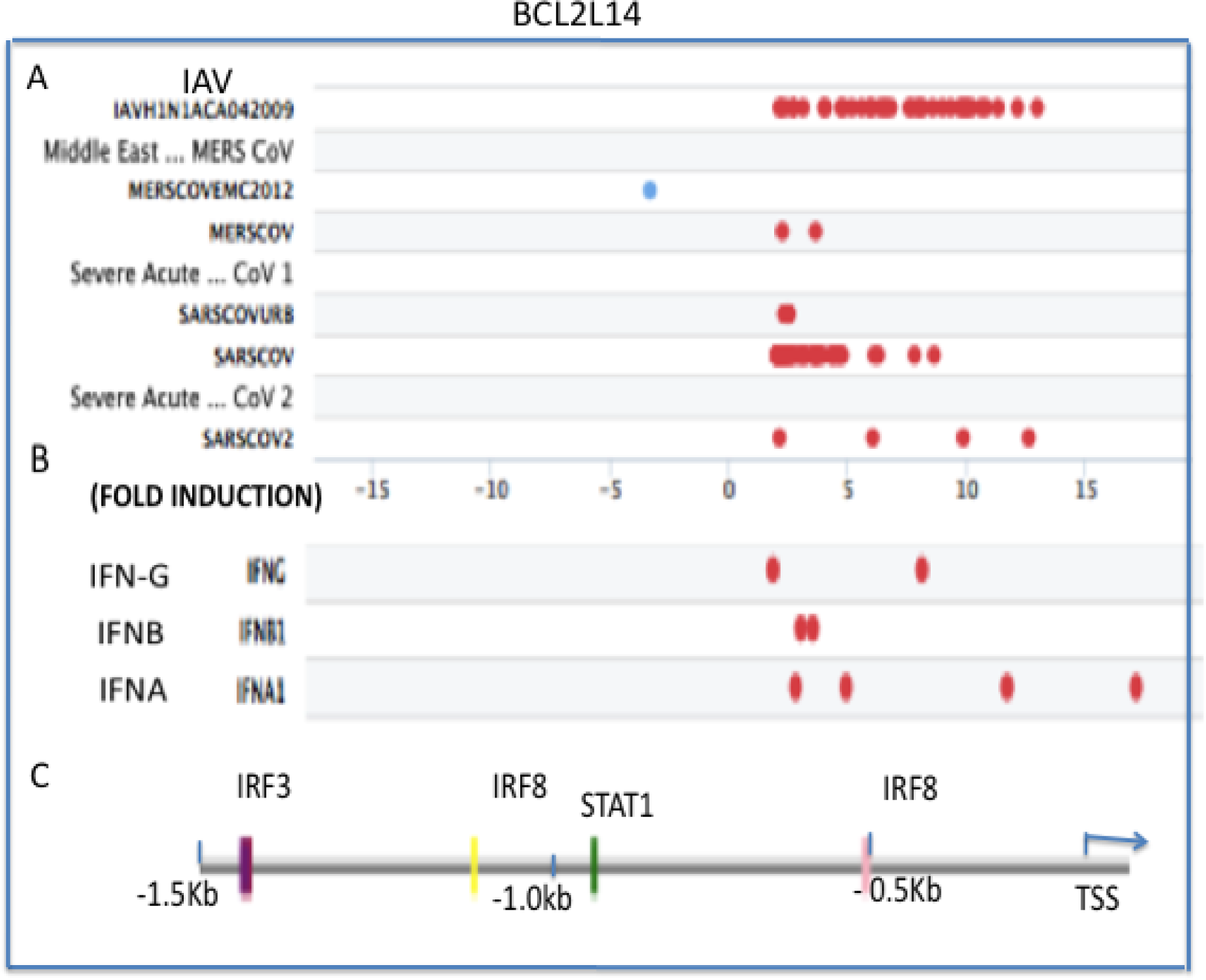
Regulation of BCL2L14 mRNA levels by respiratory viruses and interferons as represented in microarray datasets (A) Regulation of BCL2l14 mRNA levels by respiratory viruses. Induction or suppression were represented by red and blue ovals, respectively. Each oval represents a separate experiment (B) Induction of BCL2L14 by IFN−γ (IFNG), IFN-β (ΙFNB), and IFN-α (IFNA) (C) Schematic representation of the human BCL2L14 gene promoter, and the approximate location of the STAT1, IRF3, and IRF8 binding sites was shown. TSS represents the transcription start site.

### 3.7 Regulation of BCL3 and BCL6 by coronaviruses

BCL3 was originally discovered by its translocation into the immunoglobulin alpha locus in B-cell leukemia and over-expressed in some chronic lymphocytic leukemia (64). BCL3 is a member of the IKB family of proteins, associates with NF-KB (P50) homodimers, and functions as a transcriptional co- activator in inflammatory response as well as regulation of nuclear location of P50 subunit constituting an autoregulatory loop of NF-KB activation (65). BCL3 expression was induced by respiratory syncytial virus infection leading to the antagonistic effects on STAT/IRF1 and NF-KB signaling and attenuation of interleukin-8 (IL-8) expression in A549 cells (66). BCL6 is a zinc finger transcription factor required for germinal center (GC) formation, antibody maturation and functions as a repressor of genes involved in differentiation, inflammation, and apoptosis (67). Expression of a dominant-negative BCL6 in A549 lung epithelial cells stimulated chemokine gene expression in response to interleukin-4 (IL-4) and TNF– α (68). Furthermore, the amount of BCL6 bound to the promoter region of chemokine genes was decreased and STAT6 binding was enhanced in association with changes in histone acetylation and methylation markers of chromatin (68). In transgenic mice expressing BCL6, chemokine gene expression, infiltration of immune cells, and allergic inflammation were inversely correlated with the expression levels of BCL6 (68). In contrast, conditional deletion of BCL6 in neutrophils but not in monocytes or lung macrophages resulted in increased expression of apoptotic genes, neutrophil apoptosis, and reduced pulmonary inflammation in response to influenza (IAV) infection without reducing viral titers (69). Interrogation of microarray datasets revealed that BCL3 and BCL6 were regulated by inflammatory cytokines (TNF-α and IL-1β), interferons, and the steroid hormone dexamethasone (Supplementary data). Furthermore, BCL3 and BCL6 were significantly induced by coronaviruses including MERS, SARS-CoV-1, and SARS-CoV-2 (Figure 14).

**Figure 14.**
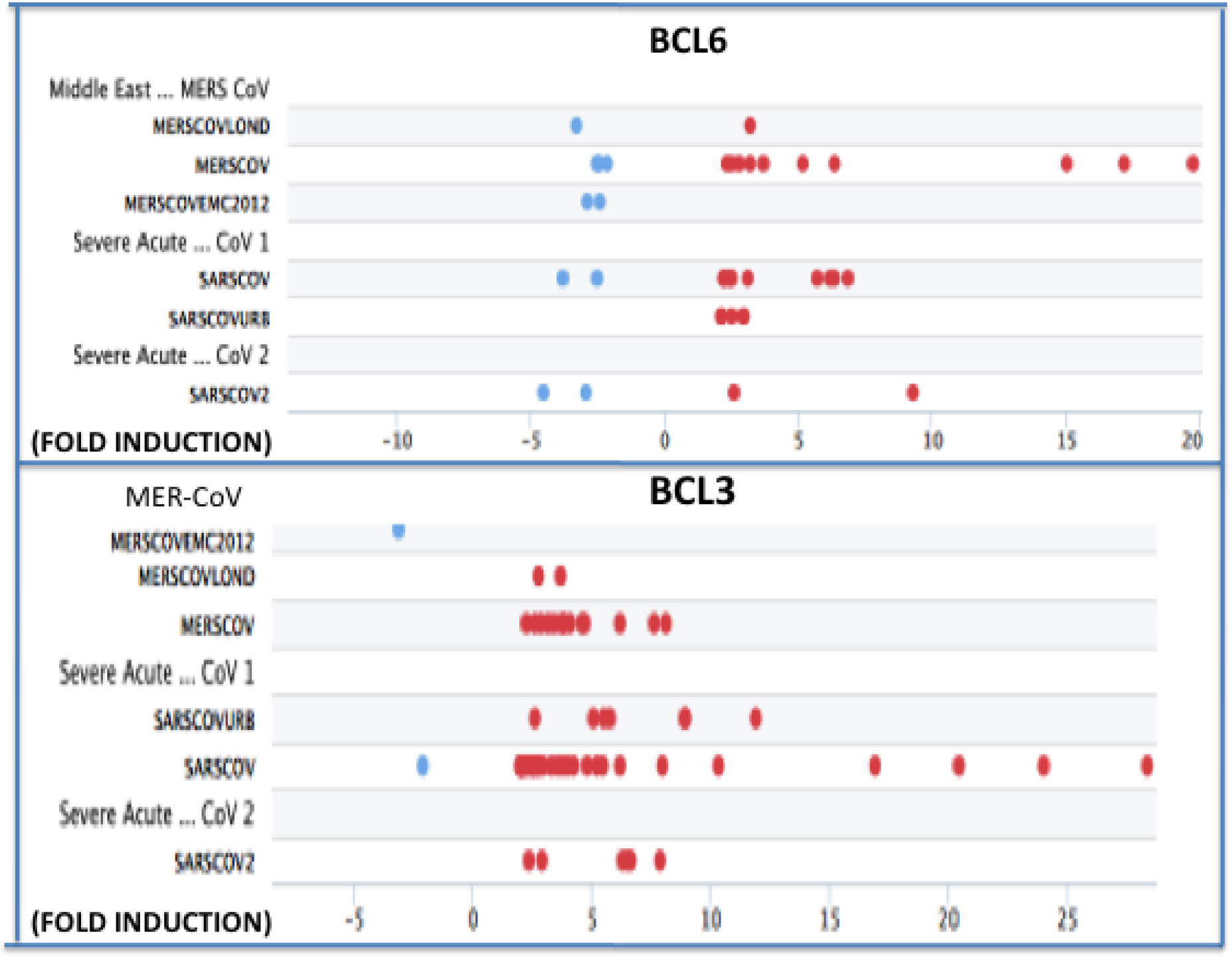
Regulation of BCL3 and BCL6 mRNA levels by respiratory viruses as represented in the microarray datasets. Induction or suppression of BCL3 or BCL6 mRNA levels were represented by red and blue ovals, respectively. Each oval represents a separate experiment.

### 3.8 Protein interaction network of BCL2A1

Protein interactions play a major role in signal transduction pathways involving post-translational modifications such as phosphorylation, stabilization, activation or repression and alteration of biological functions (30, 70). Protein interactions of BCL2A1 were visualized in the STRING database (Figure 15). These include interactions with transcription factors such as TP53, RELA, NFKB1, and apoptosis regulators such as BID, BAK1, BCL2L1, BCL2L11, and MCL1. Metascape analysis revealed that the most significant biological pathway terms or gene ontogeny (GO) terms associated with the input of the list of these gene s include apoptosis (hsa04210 in KEGG database), regulation of release of cytochrome c from mitochondria (GO0090199), and extrinsic apoptosis signaling pathway (GO0097191). It is well recognized that transcription factors TP53 and NF-KB play a major role in apoptosis in response to DNA-damaging agents and proinflammatory cytokines, respectively (71, 72). Furthermore, the BCL2A1 protein interaction list was associated with infection in DisGeNET and up-regulated genes in response to SARS-CoV-2 infection in lung epithelial cells in Metascape analysis.

**Figure 15.**
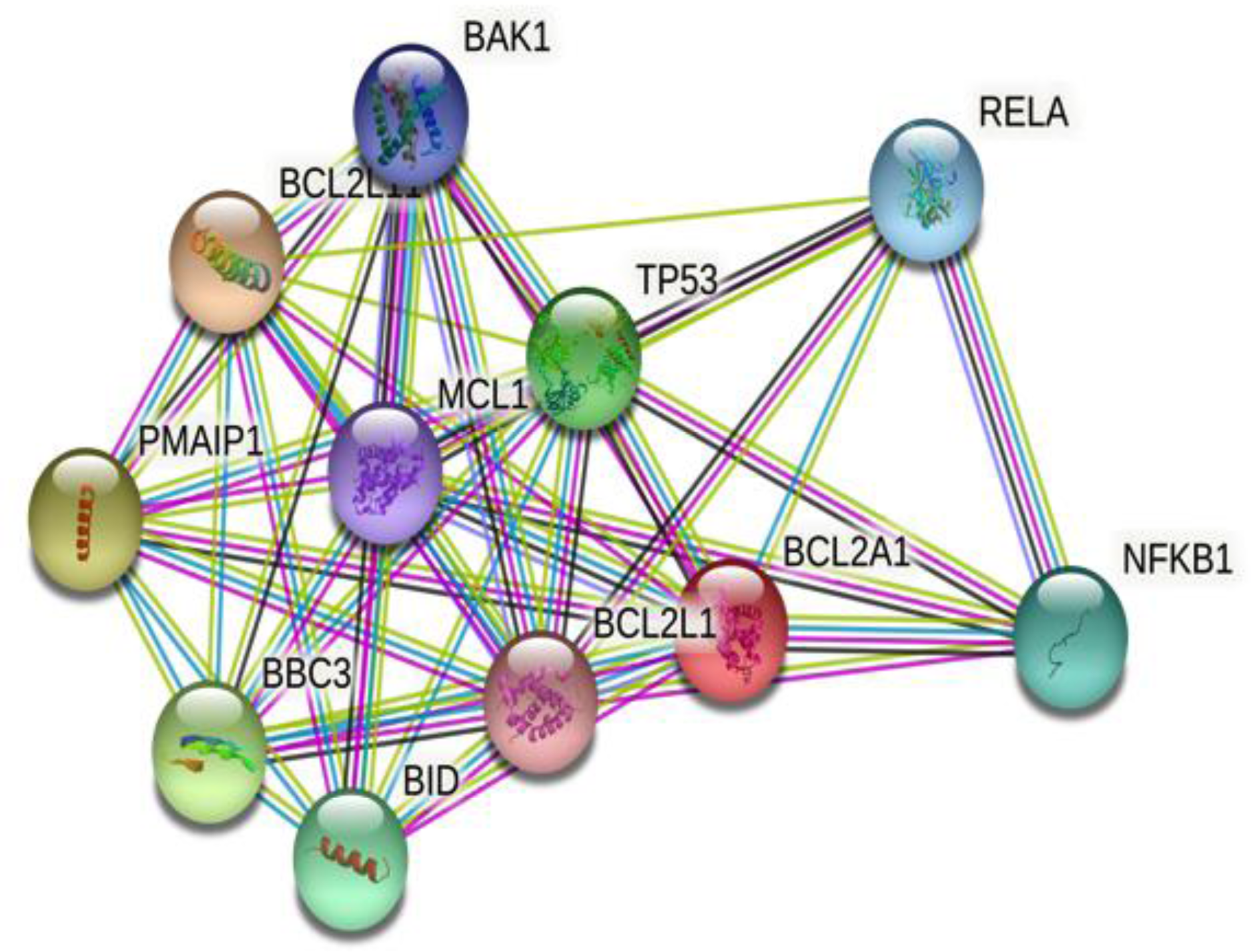
BCL2A1 and its protein-protein interaction (PPI) network. Interactions of BCL2A1 and other proteins were represented in the STRING database. Interacting proteins were indicated by ovals and protein interactions by edges, respectively.

### 3.9 Regulation of BCL2 family members in Healthy and COVID-19 patients

The overwhelming immune response to a highly pathogenic viral infection leads to an excess production of many inflammatory mediators and cellular dysfunction (47). The combination of uncontrolled inflammatory and coagulation responses results in tissue injury, deterioration of tissue and organ function, metabolic abnormalities, and potentially leading to death (13,14,28). Regulation of BCL-2 family members in host-pathogen interactions and in response to respiratory virus infection remains to be explored (73). Age-dependent higher expression of viral entry factors ACE2, TMPRSS2. and BCL-2 family members such as BCL-2, BMF, BID was reported in lung epithelial tissue samples in mice and in humans (74). Dysregulation of mitochondria and microbiota has been implicated in the pathogenesis of COVID-19 (75). Caspase 8 activation has been shown to play an important role in promoting inflammation in response to SARS-CoV-2 infection by the cleavage of the pro-IL1-β leading to the secretion of mature IL-1β leading to inflammation. Furthermore, activated caspase 8 mediated the cleavage of BID to tBID, release of the cytochrome C, caspase 3 activation and promoted mitochondrial apoptosis (20). Human bronchial epithelial cells (BEAS-2) and primary human umbilical vein endothelial cells (HUVECs) were resistant to SARS-CoV-2 infection and failed to undergo apoptosis. However, co-culture of SARS-CoV-2 infected Vero cells with BEAS-2 or HUVEC induced apoptosis in these cells without altering the permissiveness to viral infection (20) Interrogation of microarray datasets revealed that BCL2A1 and its interaction network were differentially regulated in the lung tissue samples of healthy and COVID-19 patients. RNA levels of BCL2A1,TP53, and BID were significantly increased and BCL2 L1, BCL2L11, and MCL1 levels were decreased in the lung tissue samples of COVID-19 patients in comparison with healthy controls (Figure 16A). Transcription factor mRNA like BCL3, BCL6, BCL11A were significantly increased while FOXO1 and FOXO4 involved in mitochondrial apoptosis were decreased in the lung tissue samples of COVID-19 patients in comparison with healthy controls (Figure 16B). Double-stranded RNA protein kinase or PKR (also known as EIF2AK2) was implicated in RNA-mediated apoptosis and in septic shock (24, 25). RNA expression levels of PKR were significantly increased in the lung tissue of COVID 19 patients (Figure 16B).

**Figure 16.**
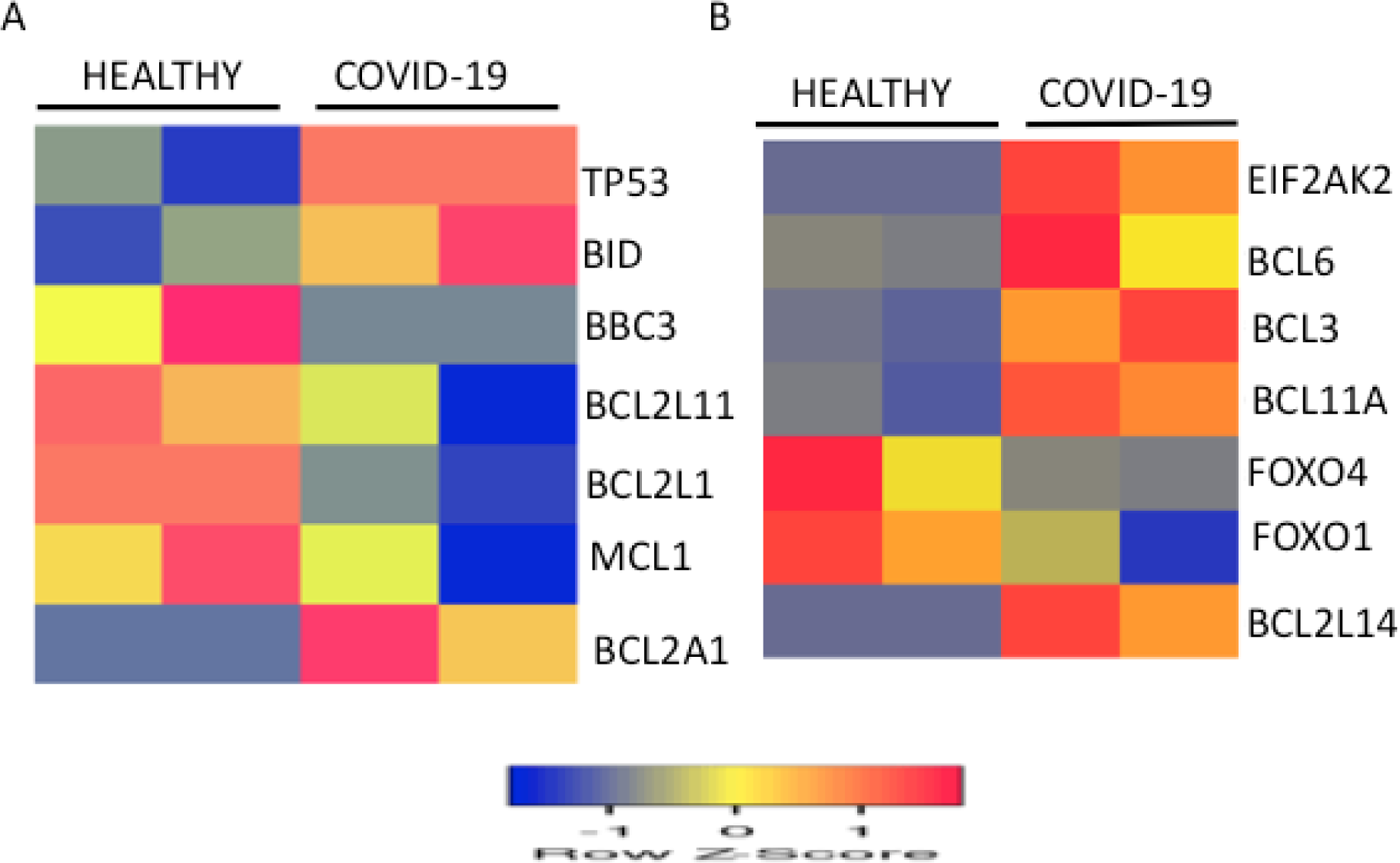
Regulation of RNA levels of selected BCL-2 family members and their interacting network in healthy controls and COVID-19 patients. (A) RNA expression levels of BCL2A and its interaction network in the healthy and COVID-19 lung samples were shown. (B) Expression levels of selected BCL-2 family transcription factors in the healthy and COVID-19 lung samples were shown. Low and high expression in healthy and COVID-19 lung biopsies were represented by blue and red, respectively. RNA expression values were normalized by the DEseq2 method and shown as a heat map. Data from two samples for each condition were shown.

## 4. CONCLUSION

Gene expression is dynamically regulated in a cell-specific manner in tissues and organs by the pathogens and extracellular ligands in eukaryotes (5, 53). Human Protein Atlas provides detailed information on gene expression at tissue and cell-specific resolution. BCL-2 family members plays a major role in immune responses such as inflammation and controlling apoptosis that are important in the clearance of respiratory viruses. Cell-specific profiling of BCL-2 family members in the human lung revealed distinct RNA expression patterns in immune cells and the AT2 cells. Interrogation of microarray datasets revealed that BCL2A1 was regulated by respiratory viruses such as SARS-CoV-2, IAV, RSV and HPIV3 and type I interferon signaling in human lung epithelial cells. Treatment of COVID-19 patients with increased oxygen therapy or hyperoxia may induce BCL2A1 expression (76). Cyclic mechanical stretching or abnormal shape of alveolar epithelial cells during ventilator-mediated lung injury may further enhance cytokine induced BCL2A1 expression in lung type II cells (54,77,78). Furthermore, additional members of the BCL-2 family such as BCL2L14, BCL3 and BCL6 were also regulated by highly pathogenic coronaviruses such as MERS, SARS-CoV1 and SARS-CoV-2 viruses. Consistent with these results, BCL members involved in transcriptional regulation and inflammation such as BCL2A1, BCL3 and BCL6 were up-regulated in the lung tissue samples of COVID-19 patients. The overall trend in transcriptomic changes in the AT2 cells favoring unbalanced cytokine responses resulting in enhanced inflammation and apoptosis contributing to lung immunopathology in the COVID-19 patients. Involvement of mitochondria in the induction of BCL2A1 via TP53, oxidative stress, and mitogen-activated protein kinase (MAPK) pathway was reported (79). BCL-2 members also regulate the metabolic functions of the mitochondria, the power houses of the cell involved in energy homeostasis (80). Transcription factor binding site data in the gene promoter regions of BCL2A1, BCL2L14, BCL3, and BCL6 suggest a complex interplay of STAT1, STAT3, IRF1, IRF7, and NF-KB transcription factors in response to extracellular signals such as TLR ligands, interferons, proinflammatory cytokines and steroid hormones (supplementary data). Transcriptomic data also revealed that the steroid hormone dexamethasone down-regulated BCL2A1and BCL3 and up-regulated BCL2L14 and BCL6 in hepatocytes (Supplementary data). Whether such regulation of BCL-2 members occurs in human lung epithelial cells or its contribution to the reported therapeutic effectiveness of corticosteroid treatment in the COVID-19 patients remains to be explored.

