## Supplementary Table 1 for "Lung Epithelial Regulation of BCL2 Related Protein A1 (BCL2A1) by Coronaviruses (SARS-CoV) and Type I Interferon Signaling"

### SUPPLEMENTARY DATA

#### BCL2A1

| **IMMGEN** | Species | Tissue | Symbol | fold change | Treatment |
|---|---|---|---|---|---|
| polyICvsVe | Mouse | AlveolarMac | Bcl2a1a | 2 | PolyIC |
| **Interferome** | | | | | |
| IFNA vs Ve | Human | Monocyte | BCL2A1 | 3.3 | IFN |
| GMCSF vs | Mouse | MDSC | Bcl2a1a | suppression | GM-CSF |
| GMCSF+IFNg | | | | | +IFNg |

**Transcriptomic data** VE= vehicle (control)

| **Experimen** | **Species** | **Tissue** | **Probe** | **Symbol** | **Fold Chang** | **P-value** | **Treatment** | |
|---|---|---|---|---|---|---|---|---|
| IL1B vs Veh | Human | hepatocyte | 205681_at | BCL2A1 | 4.08561 | 0 | IL1B | **IL-1 BETA** |
| IL1B vs Veh | Human | hepatocyte | 205681_at | BCL2A1 | 7.158266 | 0 | IL1B | |
| IL1B vs Veh | Human | hepatocyte | 205681_at | BCL2A1 | 3.069695 | 0 | IL1B | |
| IL1B vs Veh | Human | hepatocyte | 205681_at | BCL2A1 | 3.230141 | 0 | IL1B | |
| IL1B vs Veh | Human | hepatocyte | 205681_at | BCL2A1 | 2.845869 | 0 | IL1B | |
| IL1B vs Veh | Human | hepatocyte | 205681_at | BCL2A1 | 7.758253 | 0 | IL1B | |
| IL1B vs Veh | Human | hepatocyte | 205681_at | BCL2A1 | 3.124503 | 0 | IL1B | |
| IL1B vs Veh | Human | hepatocyte | 205681_at | BCL2A1 | 7.947128 | 0 | IL1B | |
| TNF vs Veh | Human | epithelium | 7990818 | BCL2A1 | 7.919604 | 0.001424 | TNF | **TNF-ALPHA** |
| TNF vs Veh | Human | hepatocyte | 205681_at | BCL2A1 | 2.236581 | 1.31E-08 | TNF | |
| TNF vs Veh | Human | hepatocyte | 205681_at | BCL2A1 | 2.869731 | 6E-10 | TNF | |
| TNF vs Veh | Human | hepatocyte | 205681_at | BCL2A1 | 2.684751 | 1.2E-09 | TNF | |
| TNF vs Veh | Human | epidermis, | ILMN_176 | BCL2A1 | 2.009518 | 0.010482 | TNF | |
| FGF19 + TN | Human | epithelium | 7990818 | BCL2A1 | 4.836208 | 0.003608 | FGF19,TNF | |
| LPS vs Veh | Norway Ra | whole liver | 1368482_a | Bcl2a1 | 6.153416 | 0.000231 | LPS | **TLR** |
| LPS vs Veh | Human | hepatocyte | 205681_at | BCL2A1 | 6.314044 | 0 | LPS | |
| LPS vs Veh | Norway Ra | whole liver | 1368482_a | Bcl2a1 | 5.880627 | 0.000311 | LPS | |
| LPS vs Veh | Norway Ra | whole liver | 1368482_a | Bcl2a1 | 2.689758 | 0.031984 | LPS | |
| LPS vs Veh | Human | hepatocyte | 205681_at | BCL2A1 | 2.838135 | 6E-10 | LPS | |
| LPS vs Veh | Norway Ra | whole liver | 1368482_a | Bcl2a1 | 6.090048 | 0.000247 | LPS | |
| LPS vs Veh | Human | hepatocyte | 205681_at | BCL2A1 | 4.861197 | 0 | LPS | |
| LPS vs Veh | Norway Ra | whole liver | 1368482_a | Bcl2a1 | 6.043804 | 0.00026 | LPS | |
| LPS vs Veh | Human | hepatocyte | 205681_at | BCL2A1 | 4.242685 | 0 | LPS | |
| P3CSK4 vs | Human | myeloid lin | BCL2A1\|96 | BCL2A1 | 2.782493 | 5.77E-08 | P3CSK4 | TLR |
| P3CSK4 vs | Human | myeloid lin | BCL2A1\|96 | BCL2A1 | 5.574397 | 0 | P3CSK4 | |
| 125DVD3 v | Human | myeloid lin | 205681_at | BCL2A1 | 8.510467 | 2.81E-05 | 125DVD3 | **STEROID** |
| 125DVD3 v | Human | smooth mu | BCL2A1\|96 | BCL2A1 | 2.520448 | 0.0012 | 125DVD3 | VITAM D3 |
| 125DVD3 v | Human | smooth mu | BCL2A1\|96 | BCL2A1 | 7.105617 | 0.0006 | 125DVD3 | |
| | | | | **DOWN-REGULATED** | | | | |
| TGFB2 vs V | Human | other, fibro | 205681_at | BCL2A1 | -2.42428 | 0.007574 | TGFB2 | **TGF BETA** |
| TGFB1 vs V | Human | other, fibro | 205681_at | BCL2A1 | -2.52269 | 0.006081 | TGFB1 | |
| TGFB1 vs V | Human | hepatocyte | 205681_at | BCL2A1 | -4.42311 | 6E-10 | TGFB1 | |
| TGFB1 vs V | Human | hepatocyte | 205681_at | BCL2A1 | -4.59256 | 4E-10 | TGFB1 | |
| TGFB1 vs V | Human | hepatocyte | 205681_at | BCL2A1 | -4.94485 | 2E-10 | TGFB1 | |
| DEX vs Veh | Norway Ra | whole liver | 1368482_a | Bcl2a1 | -2.91714 | 2.52E-05 | DEX | **STEROID** |
| DEX vs Veh | Norway Ra | whole liver | 1368482_a | Bcl2a1 | -3.78112 | 6.78E-07 | DEX | DEXAMETH |
| DEX vs Veh | Norway Ra | whole liver | 1368482_a | Bcl2a1 | -2.63061 | 0.000103 | DEX | |
| DEX vs Veh | Norway Ra | whole liver | 1368482_a | Bcl2a1 | -2.86176 | 3.28E-05 | DEX | |

| | | | | | | | |
|---|---|---|---|---|---|---|---|
| DEX vs Veh | Norway Ra | whole liver | A_42_P518 | Bcl2a1 | -3.92259 | 0.000206 | DEX |
| DEX vs Veh | Norway Ra | whole liver | A_42_P518 | Bcl2a1 | -4.57337 | 8.82E-05 | DEX |
| DEX vs Veh | Norway Ra | whole liver | 1368482_a | Bcl2a1 | -2.37092 | 0.000413 | DEX |
| DEX vs Veh | Norway Ra | whole liver | 1368482_a | Bcl2a1 | -2.00596 | 0.003401 | DEX |
| DEX vs Veh | Norway Ra | whole liver | 1368482_a | Bcl2a1 | -2.91024 | 2.6E-05 | DEX |
| DEX vs Veh | Norway Ra | whole liver | 1368482_a | Bcl2a1 | -4.36257 | 9.24E-08 | DEX |
| DEX vs Veh | Norway Ra | whole liver | 1368482_a | Bcl2a1 | -3.6783 | 9.97E-07 | DEX |
| DEX vs Veh | Human | epithelium | BCL2A1 | BCL2A1 | -3.61125 | 1E-10 | DEX |
| DEX vs Veh | Human | hepatocyte | 205681_at | BCL2A1 | -2.68814 | 4.22E-06 | DEX |
| DEX vs Veh | Norway Ra | whole liver | 1368482_a | Bcl2a1 | -2.24437 | 0.000368 | DEX |

**BCL2L14**

**INTERFEROME DATA**

| DATASET ID | FOLD INDUCTION | TREATMENT TIME | TYPE OF INTERFERON | PROBESET | | PROMOTER TF DATA STAT1,STAT3,NF-KB |
|---|---|---|---|---|---|---|
| 47 | 3.9 | 5 HR | Type I | 221241_s_at | | |
| 301 | 7 | 6 HR | Type I | 7953993 | | |

**TRANSCRIPTOMIC DATA**

| Experimen | Species | Tissue | Probe | Symbol | Fold Chang | P-value | Treatment |
|---|---|---|---|---|---|---|---|
| IFNA1 vs V | Human | hepatocyte | 221241_s_ | BCL2L14 | 23.89382 | 0 | IFNA1 |
| IFNB1 vs C | Human | epithelium | BCL2L14 | BCL2L14 | 4.917129 | 0.001162 | IFNB1 |
| IFNA1 vs V | Human | hepatocyte | 221241_s_ | BCL2L14 | 4.103503 | 8E-09 | IFNA1 |
| IFNG vs Ve | Human | myeloid lir | BCL2L14\|9 | BCL2L14 | 9.694558 | 0 | IFNG |
| IFNA1 vs V | Human | hepatocyte | 221241_s_ | BCL2L14 | 13.41221 | 0 | IFNA1 |
| IFNG vs Ve | Human | epidermis, | 221241_s_ | BCL2L14 | 3.112359 | 0.010527 | IFNG |
| IFNA1 vs V | Human | hepatocyte | 221241_s_ | BCL2L14 | 19.09894 | 0 | IFNA1 |
| IFNB1 vs C | Human | epithelium | BCL2L14 | BCL2L14 | 4.367666 | 0.005981 | IFNB1 |
| IFNA1 vs V | Human | hepatocyte | 221241_s_ | BCL2L14 | 6.341885 | 2E-10 | IFNA1 |
| TGFB2 vs V | Human | other, fibro | 221241_s_ | BCL2L14 | -3.8638 | 0.021441 | TGFB2 |
| TGFB2 vs V | Human | other, fibro | 1552841_s | BCL2L14 | -2.97559 | 0.002472 | TGFB2 |
| 17BE2 vs V | Human | epithelium | NM_13872 | BCL2L14 | -8.69614 | 0.002503 | 17BE2 |
| 17BE2 vs V | Human | epithelium | NM_03076 | BCL2L14 | -8.69614 | 0.002503 | 17BE2 |
| 17BE2 vs V | Human | epithelium | NM_13872 | BCL2L14 | -8.69614 | 0.002503 | 17BE2 |
| DHT vs Veh | Human | epithelium | 1552841_s | BCL2L14 | -4.45158 | 0.025096 | DHT |
| DEX vs Veh | Human | lymphoid l | 221241_s_ | BCL2L14 | 3.300185 | 0.041228 | DEX |
| DEX vs Veh | Human | eye, lens, e | 221241_s_ | BCL2L14 | 3.397556 | 0.022652 | DEX |
| RU486 + D | Human | epithelium | 233464_at | BCL2L14 | 4.506866 | 0.036092 | DEX,RU486 |
| RIP140/NR | Human | pluripoten | 1552841_s | BCL2L14 | 7.922585 | 0.000801 | ATRA |
| RIP140/NR | Human | pluripoten | 1552841_s | BCL2L14 | 7.922585 | 0.000801 | ATRA |
| ATRA vs Ve | House Mou | pluripoten | Bcl2l14\|10 | Bcl2l14 | -2.88126 | 0.00005 | ATRA |
| ATRA vs Ve | Human | pluripoten | 1552841_s | BCL2L14 | 2.76016 | 0.040375 | ATRA |

BCL3

**INTERFEROME DATA**

| DATASET ID | FOLD INDUCTION | TREATMENT TIME | TYPE OF INTERFERON | PROBESET | | PROMOTER TF DATA STAT1,STAT3,NF-KB |
|---|---|---|---|---|---|---|
| 306 | 2.5 | 1.5HR | TYPE I | 204908_s_at | | |
| 282 | 4.3 | 10 HR | TYPE I | ILMN_2749717 | | |

**TRANSCRIPTOME DATA**

| Experimen | Species | Tissue | Probe | Symbol | Fold Chang | P-value | TREATMENT |
|---|---|---|---|---|---|---|---|
| TNF vs Veh | Human | epidermis, | ILMN_171( | BCL3 | 4.288766 | 8.12E-07 | TNF |
| LPS vs Veh | Norway Ra | whole liver | 1398482_a | Bcl3 | 11.62403 | 7.28E-08 | LPS |
| LPS vs Veh | Norway Ra | whole liver | 1380414_a | Bcl3 | 2.730324 | 0.000159 | LPS |
| LPS vs Veh | Norway Ra | whole liver | 1380414_a | Bcl3 | 2.492361 | 0.000484 | LPS |
| LPS vs Veh | Norway Ra | whole liver | 1398482_a | Bcl3 | 2.597225 | 0.012396 | LPS |
| LPS vs Veh | Norway Ra | whole liver | 1380414_a | Bcl3 | 2.487383 | 0.000496 | LPS |
| LPS vs Veh | Norway Ra | whole liver | 1385627_a | Bcl3 | 2.052888 | 0.006553 | LPS |
| IL1B vs Veh | Human | hepatocyte | 204908_s_ | BCL3 | 2.107252 | 2.5E-09 | IL1B |
| LPS vs Vehl | House Mou | myeloid lin | 1418133_a | Bcl3 | 4.858643 | 1.62E-08 | LPS |
| TNF vs Veh | Human | epidermis, | ILMN_171( | BCL3 | 5.469214 | 9.85E-08 | TNF |
| IL1B vs Veh | Human | hepatocyte | 204908_s_ | BCL3 | 2.19696 | 1.1E-09 | IL1B |
| LPS vs Veh | Norway Ra | whole liver | 1385627_a | Bcl3 | 4.576919 | 5.39E-07 | LPS |
| LPS vs Veh | Norway Ra | whole liver | 1398482_a | Bcl3 | 7.712182 | 2.25E-06 | LPS |
| TNF vs Veh | Human | epidermis, | ILMN_171( | BCL3 | 3.268819 | 0.000011 | TNF |
| LPS vs Veh | House Mou | whole liver | 1418133_a | Bcl3 | 2.153395 | 0.001591 | LPS |
| LPS vs Veh | Norway Ra | whole liver | 1398482_a | Bcl3 | 2.427828 | 0.019411 | LPS |
| IL6 vs Veh | Human | hepatocyte | 204908_s_ | BCL3 | 2.328255 | 1.46E-08 | IL6 |
| LPS vs Veh | Norway Ra | whole liver | 1398482_a | Bcl3 | 2.562114 | 0.013588 | LPS |
| IL1B vs Veh | Human | hepatocyte | 204908_s_ | BCL3 | 2.198495 | 1.1E-09 | IL1B |
| LPS vs Veh | Norway Ra | whole liver | 1385627_a | Bcl3 | 2.056721 | 0.006429 | LPS |
| TGFB1 vs V | House Mou | epithelium | 1418133_a | Bcl3 | 2.304235 | 2.47E-05 | TGFB1 |
| LPS vs Veh | Norway Ra | whole liver | 1380414_a | Bcl3 | 2.186874 | 0.002252 | LPS |
| LPS vs Veh | Norway Ra | whole liver | 1380414_a | Bcl3 | 2.118609 | 0.003225 | LPS |
| TNF + EVs v | Human | myeloid lin | A_23_P46( | BCL3 | 3.379305 | 0.000185 | TNF |
| LPS vs Veh | Norway Ra | whole liver | 1385627_a | Bcl3 | 2.063246 | 0.006222 | LPS |
| LPS vs Veh | House Mou | ventricle, v | 1418133_a | Bcl3 | 3.070556 | 4.29E-06 | LPS |
| IL6 vs Veh | House Mou | hepatocyte | 1418133_a | Bcl3 | 2.192341 | 2.14E-05 | IL6 |
| TNF vs Veh | Norway Ra | whole liver | 1398482_a | Bcl3 | 3.380392 | 0.000413 | TNF |
| LPS vs Veh | Norway Ra | whole liver | 1398482_a | Bcl3 | 2.472971 | 0.017202 | LPS |
| LPS vs Veh | Norway Ra | whole liver | 1380414_a | Bcl3 | 3.662326 | 3.85E-06 | LPS |
| IL6 vs Veh | Human | hepatocyte | 204908_s_ | BCL3 | 2.084584 | 1.15E-07 | IL6 |
| LPS vs Veh | Norway Ra | whole liver | 1385627_a | Bcl3 | 3.058034 | 0.000073 | LPS |
| TNF vs Veh | Norway Ra | whole liver | 1398482_a | Bcl3 | 2.400989 | 0.008244 | TNF |
| TNF vs Veh | Human | epidermis, | ILMN_171( | BCL3 | 3.187529 | 1.42E-05 | TNF |
| LPS vs Veh | Norway Ra | whole liver | 1380414_a | Bcl3 | 3.678189 | 3.64E-06 | LPS |
| IL6 vs Veh | House Mou | whole liver | 1418133_a | Bcl3 | 2.095324 | 0.027348 | IL6 |
| LPS vs Veh | House Mou | ventricle, v | 1418133_a | Bcl3 | 4.419092 | 7.17E-08 | LPS |
| LPS vs Veh | House Mou | whole liver | 1418133_a | Bcl3 | 5.445285 | 1.72E-06 | LPS |

| | | | | | | | |
|---|---|---|---|---|---|---|---|
| LPS vs Veh | Norway Ra | whole liver | 1398482_a | Bcl3 | 11.63733 | 7.21E-08 | LPS |
| LPS vs Veh | Norway Ra | whole liver | 1385627_a | Bcl3 | 3.381128 | 2.18E-05 | LPS |
| TNF vs Veh | Norway Ra | whole liver | 1398482_a | Bcl3 | 2.223592 | 0.015142 | TNF |
| IL6 vs Veh | Human | hepatocyte | 204908_s_ | BCL3 | 2.008681 | 2.41E-07 | IL6 |
| IL1B vs Veh | Human | hepatocyte | 204908_s_ | BCL3 | 2.058651 | 4E-09 | IL1B |
| TNF vs Veh | Human | hepatocyte | 204908_s_ | BCL3 | 2.022049 | 2E-07 | TNF |
| IL6 vs Veh | Human | hepatocyte | 204908_s_ | BCL3 | 2.099584 | 1E-07 | IL6 |
| LPS vs Veh | Norway Ra | whole liver | 1385627_a | Bcl3 | 2.096701 | 0.005265 | LPS |
| LPS vs Veh | Norway Ra | whole liver | 1385627_a | Bcl3 | 4.222079 | 1.44E-06 | LPS |
| LPS vs Veh | Norway Ra | whole liver | 1398482_a | Bcl3 | 5.245665 | 5.65E-05 | LPS |
| IL6 vs Veh | Human | hepatocyte | 204908_s_ | BCL3 | 2.3857 | 9.5E-09 | IL6 |
| IL6 vs Veh | Human | hepatocyte | 204907_s_ | BCL3 | 2.063463 | 1.67E-06 | IL6 |
| LPS vs Veh | Human | hepatocyte | 204908_s_ | BCL3 | 2.098664 | 7.7E-08 | LPS |
| TNF vs Veh | Human | myeloid lin | A_23_P46 | BCL3 | 3.209034 | 0.000264 | TNF |
| P3CSK4 vs | Human | myeloid lin | BCL3\|9606 | BCL3 | 3.068522 | 0.000251 | P3CSK4 |
| DEX vs Veh | Human | osteoblast | 204908_s_ | BCL3 | -2.37076 | 1.6E-09 | DEX |
| DEX vs Veh | Human | osteoblast | A_23_P46 | BCL3 | -5.34867 | 3E-10 | DEX |
| DEX vs Veh | Human | osteoblast | A_23_P46 | BCL3 | -5.18254 | 4E-10 | DEX |
| DEX vs Veh | House Mou | whole liver | A_51_P13 | Bcl3 | -2.97125 | 0.016511 | DEX |
| DEX vs Veh | Human | epithelium | ILMN_171 | BCL3 | -2.4007 | 0.000384 | DEX |
| DEX vs Veh | Human | osteoblast | A_23_P46 | BCL3 | -5.09418 | 5E-10 | DEX |
| LIF vs Veh | House Mou | pituitary, c | 1418133_a | Bcl3 | 6.758471 | 1.75E-07 | LIF |
| LIF vs Veh | House Mou | pituitary, c | 1418133_a | Bcl3 | 5.217156 | 5.97E-07 | LIF |
| Dex + LIF v | House Mou | pituitary, c | 1418133_a | Bcl3 | 5.543989 | 4.41E-07 | DEX,LIF |
| Dex + LIF v | House Mou | pituitary, c | 1418133_a | Bcl3 | 5.261682 | 5.72E-07 | DEX,LIF |
| 125DVD3 v | Human | myeloid lin | 204907_s_ | BCL3 | 2.29902 | 0.007805 | 125DVD3 |
| 125DVD3 v | Human | epithelium | 204908_s_ | BCL3 | 2.270671 | 3.73E-05 | 125DVD3 |
| ATRA vs Ve | Human | myeloid lin | 204907_s_ | BCL3 | 7.855398 | 0.001219 | ATRA |
| ATRA vs Ve | House Mou | pluripoten | Bcl3\|1009 | Bcl3 | -2.75604 | 1.61E-06 | ATRA |
| ATRA vs Ve | House Mou | pluripoten | Bcl3\|1009 | Bcl3 | -6.31595 | 0.00005 | ATRA |

**BCL6**

**INTERFEROME DATA**

| DATASET ID | FOLD INDUCTION | TREATMENT TIME | TYPE OF INTERFERON | PROBESET | | PROMOTER TF DATA STAT1,STAT3,NF-KB,IRF |
|---|---|---|---|---|---|---|
| 26 | 3.1 | 2 HR | TYPE I | 203140_at | | |
| 377 | 2 | 4 HR | TYPE I | 1421818_at | | |

**TRANSCRIPTOMIC DATA**

| Experimen | Species | Tissue | Probe | Symbol | Fold Chang | P-value | TREATMENT |
|---|---|---|---|---|---|---|---|
| IFNA1 vs V | Human | hepatocyte | 215990_s_ | BCL6 | 2.574703 | 4.29E-08 | IFNA1 |
| IFNA1 vs V | Human | hepatocyte | 215990_s_ | BCL6 | 2.743578 | 1.84E-08 | IFNA1 |
| IFNA1 vs V | Human | hepatocyte | 203140_at | BCL6 | 3.461544 | 0 | IFNA1 |
| IFNA1 vs V | Human | hepatocyte | 203140_at | BCL6 | 2.718479 | 0 | IFNA1 |
| LPS vs Veh | House Mo | whole liver | 1421818_a | Bcl6 | 2.967737 | 0.045892 | LPS |
| LPS vs Veh | Norway Ra | whole liver | 1379368_a | Bcl6 | 5.934082 | 0.00498 | LPS |
| IL6 vs Veh | Human | epithelium | 215990_s_ | BCL6 | -2.0305 | 0.03475 | IL6 |
| IL1B vs Veh | Human | hepatocyte | 203140_at | BCL6 | 2.071135 | 0 | IL1B |
| TGFB1 vs V | House Mo | bone marr | A_52_P16 | Bcl6 | 2.066521 | 1.14E-05 | TGFB1 |
| TGFB1 vs V | House Mo | epithelium | 1421818_a | Bcl6 | 2.04782 | 0.000158 | TGFB1 |
| TGFB1 vs V | House Mo | epithelium | 1421818_a | Bcl6 | 2.262007 | 7.49E-05 | TGFB1 |
| TGFB1 vs V | House Mo | epithelium | 1421818_a | Bcl6 | 2.440173 | 1.92E-05 | TGFB1 |
| IL6 vs Veh | Human | hepatocyte | 203140_at | BCL6 | 3.390614 | 0 | IL6 |
| IL6 vs Veh | Human | hepatocyte | 203140_at | BCL6 | 2.750324 | 0 | IL6 |
| IL6 vs Veh | Human | epithelium | 203140_at | BCL6 | -2.6132 | 0.00067 | IL6 |
| LPS vs Veh | Human | hepatocyte | 203140_at | BCL6 | 2.331652 | 2E-10 | LPS |
| LPS vs Veh | House Mo | whole liver | 1421818_a | Bcl6 | 3.644268 | 6.39E-06 | LPS |
| DEX vs Veh | House Mo | other, neu | 1450381_a | Bcl6 | 2.13359 | 1.13E-07 | DEX |
| DEX vs Veh | House Mo | other, neu | 1421818_a | Bcl6 | 3.623291 | 3E-10 | DEX |
| LPS vs Veh | Norway Ra | whole liver | 1379368_a | Bcl6 | 3.489737 | 0.042668 | LPS |
| IL6 vs Veh | Human | hepatocyte | 215990_s_ | BCL6 | 2.914916 | 0.000014 | IL6 |
| TNF vs Veh | Human | epidermis, | ILMN_173 | BCL6 | 3.080802 | 7.62E-05 | TNF |
| IL1B vs Veh | Human | hepatocyte | 203140_at | BCL6 | 2.161568 | 0 | IL1B |
| LPS vs Veh | Norway Ra | whole liver | 1379368_a | Bcl6 | 6.53881 | 0.003238 | LPS |
| TNF vs Veh | Human | epithelium | 8092691 | BCL6 | 2.119827 | 0.000117 | TNF |
| LPS vs Vehl | House Mo | myeloid lir | 1450381_a | Bcl6 | 2.548232 | 1.59E-07 | LPS |
| IL6 vs Veh | Human | hepatocyte | 228758_at | BCL6 | 2.060861 | 5.43E-07 | IL6 |
| IL6 vs Veh | Human | epithelium | 215990_s_ | BCL6 | -3.46765 | 0.002133 | IL6 |
| LPS vs Veh | House Mo | whole liver | 1421818_a | Bcl6 | 3.233368 | 2.94E-05 | LPS |
| IL6 vs Veh | Human | hepatocyte | 203140_at | BCL6 | 3.476582 | 0 | IL6 |
| IL6 vs Veh | Human | hepatocyte | 215990_s_ | BCL6 | 2.736582 | 2.87E-05 | IL6 |
| IL1B vs Veh | Human | hepatocyte | 203140_at | BCL6 | 2.022742 | 0 | IL1B |
| IL1B vs Veh | Human | hepatocyte | 203140_at | BCL6 | 2.128554 | 0 | IL1B |
| IL1B vs Veh | Human | hepatocyte | 215990_s_ | BCL6 | 2.043159 | 3.07E-07 | IL1B |
| DEX vs Veh | Norway Ra | whole liver | A_44_P10 | Bcl6 | 4.522341 | 0.01273 | DEX |
| DEX vs Veh | Norway Ra | whole liver | A_44_P10 | Bcl6 | 6.927343 | 0.003049 | DEX |
| DEX vs Veh | Norway Ra | whole liver | 1379368_a | Bcl6 | 7.664973 | 3.73E-05 | DEX |

| | | | | | | | |
|---|---|---|---|---|---|---|---|
| DEX vs Veh | Norway Ra | whole liver | 1379368_a | Bcl6 | 5.422809 | 0.000403 | DEX |
| DEX vs Veh | Norway Ra | whole liver | 1379368_a | Bcl6 | 4.593792 | 0.001203 | DEX |
| DEX vs Veh | House Mou | cerebrum, | 1421818_a | Bcl6 | 2.499747 | 9E-10 | DEX |
| DEX vs Veh | Human | epithelium | ILMN_173 | BCL6 | 2.143576 | 1E-10 | DEX |
| ATRA vs Ve | Human | pluripoten | 228758_at | BCL6 | 5.541184 | 2.02E-06 | ATRA |
| RIP140/NR | Human | pluripoten | 228758_at | BCL6 | 4.138849 | 1.01E-05 | ATRA |
| RIP140/NR | Human | pluripoten | 228758_at | BCL6 | 4.138849 | 1.01E-05 | ATRA |
| RIP140/NR | Human | pluripoten | 203140_at | BCL6 | 3.585089 | 2E-05 | ATRA |
| RIP140/NR | Human | pluripoten | 203140_at | BCL6 | 3.585089 | 2E-05 | ATRA |
| ATRA vs Ve | House Mou | pluripoten | Bcl6\|1009 | Bcl6 | 2.656093 | 0.00001 | ATRA |
| ATRA vs Ve | Human | pluripoten | 215990_s_ | BCL6 | 6.052565 | 0.005373 | ATRA |
| ATRA vs Ve | Human | pluripoten | 228758_at | BCL6 | 3.213533 | 5.09E-05 | ATRA |
| ATRA vs Ve | Human | pluripoten | 203140_at | BCL6 | 3.690208 | 1.66E-05 | ATRA |
| 125DVD3 v | Human | myeloid lir | 215990_s_ | BCL6 | 3.26518 | 0.00438 | 125DVD3 |
| 125DVD3 v | Human | myeloid lir | 8092691 | BCL6 | 2.143376 | 1.28E-05 | 125DVD3 |
| 125DVD3 v | Human | myeloid lir | 203140_at | BCL6 | 13.29916 | 1.06E-06 | 125DVD3 |
